# Longer internal exons tend to have more tandem repeats and more frequently experience insertions and deletions that are mostly in intrinsically disordered regions of the encoded proteins

**DOI:** 10.1101/2024.11.03.621752

**Authors:** Keiichi Homma, Hiroto Anbo, Motonori Ota, Satoshi Fukuchi

## Abstract

Insertions and deletions (indels) in eukaryotic proteins are known to preferentially encode intrinsically disordered regions (IDRs), protein regions that by themselves do not form unique three-dimensional structures. As a previous investigation showed that long internal exons tend to encode IDRs in eukaryotes in general, we thought it worthwhile to analyze how indels alter internal exons and affect IDRs of the encoded proteins. For consideration of evolutionary roles indels play, we decided to select indels commonly observed in all variants (“fixed” indels) since indels in minor variants may represent transient aberrations in splicing. Here, by comparison of orthologous variants of closely related species together with those of outgroups, we identified fixed indels in the internal exons in four mammals and two flies. The fixed indels are nearly always nonframeshifting, short, and mostly encode IDRs. On average 51% of inserted and 40% of deleted residues are attributable to alterations in tandem repeats. Deletion tends to occur more frequently than insertion does and indels are generally more prevalent in long internal exons. Tandem repeats occur preferentially in long internal exons, indicating that their alterations account for the high frequency of indels in long internal exons. Also, since tandem repeats mostly encode IDRs, this finding at least partially explains the high incidence of IDRs in long internal exons. We propose that long internal exons had been produced in early eukaryotes mainly by repeat expansion that added IDRs to the encoded proteins but are experiencing frequent indels by alterations in tandem repeats.

## Introduction

Only a small fraction of prokaryotic proteins (residue-wise 2.0% of archaean and 4.2% of eubacterial proteins) consist of intrinsically disordered regions (IDRs) that by themselves do not form unique three-dimensional structures, whereas 33.0% of residues in eukaryotic proteins are comprised of IDRs (Ward et al. 2004). IDRs in eukaryotic proteins participate in binding to other molecules and are thereby involved in important functions such as gene transcription and signaling (Anbo et al. 2019). Two mechanisms of *de novo* IDR generation are expansion of repetitive DNA sequences (Tompa 2003) and exonization of introns (Kondrashov et al. 2003; Sorek 2007; Marquez et al. 2015), but to our knowledge their proportional shares have not been quantified. It is of interest to elucidate how IDRs had been acquired in the evolutionary process from the first eukaryotic common ancestor (FECA) to the last eukaryotic common ancestor (LECA).

We previously reported the general eukaryotic tendency of long internal exons to encode IDRs and proposed that long internal exons in the LECA had resulted from short exons by addition of IDR-encoding nucleotides (Fukuchi et al. 2023). As indels play important roles in exon length alterations and mostly encode IDRs (Light et al. 2013A, B; Khan et al. 2015), indel analyses may reveal mechanisms of exon length alterations and thereby explain how the preferential encoding of IDRs by long internal exons came about.

Indels in human genomes cause as much variation as small nucleotide polymorphisms (Mullaney et al. 2010) and one way to identify them in coding sequences is to find indels in genomes and select those in coding sequences (Anzai et al. 2003; Mills et al. 2006; Wetterbom et al. 2006; Fan et al. 2007; Khan et al. 2015; Lin et al. 2017). As coding sequences vary from one alternative splicing variant to another, however, different sets of coding sequences result in different indel identifications. Another method to select indels in proteins is to compare the amino acids sequences of orthologous proteins either by sequence alignments (Benner et al. 1993; Light et al. 2013A, B) or by structural alignments (Pascarella et al. 1992; Qian and Goldstein 2001). Since indels thereby identified are dependent on the selection of orthologous proteins, splicing variants need to be considered, and structural alignments cannot be made to stretches containing IDRs, which is a relevant issue because, as stated above, indels frequently encode IDRs (Light et al. 2013A, B; Khan et al. 2015). As the increasing availability of variant sequences makes it possible to identify indels shared by all variants, we conducted sequence alignments of all available variants, selected commonly observed indels, and called them “fixed” indels.

Fixed indels can result from constitutive changes in splicing and genomic mutations. Changes in splicing that give rise to insertions can be regarded as intron to exon conversions (exonizations), while those resulting in deletions can be considered as exon to intron conversions (intronizations). Indels by genomic mutations have three plausible causes: DNA slippage, DNA damage followed by imperfect repair mostly by means of homologous repair pathways (Redelings et al. 2024), and transposable elements, which constitute approximately 4% of human protein-coding genes (Nekrutenko and Li 2001). DNA slippage frequently occurs in short tandem repeats that in most cases encode IDRs (Simon and Hancock 2009, Levinson and Gutman 1987). Proteins in all kingdoms of life have tandem repeats, but prokaryotic proteins tend to have tandem repeats slightly less often than eukaryotic proteins do (Delucchi et al. 2020).

Since splicing of exons longer than 300 nt is inhibited unless flanked by short introns (Robberson et al. 1990; Chen and Chasin 1994; Sterner et al. 1996), many exons that became excessively long may not be efficiently spliced. On the other hand, analyses of RNA-Seq data demonstrated that introns shorter than 70 nt are not completely spliced out (Abebrese et al. 2017). Exon definition was proposed to explain the splicing mechanism in vertebrates in which short exons separated by long (>250 bp) introns predominate (Robberson et al. 1990; Berget 1994; De Conti et al. 2012), while intron definition was postulated to account for the splicing of long exons with short introns prevalent in lower eukaryotes (Lang and Spritz 1983; Berget 1994; De Conti et al. 2012). Recent research disclosed that the same spliceosome assembled on introns provides the mechanisms for both intron and exon definition in *Saccharomyces cerevisiae* (Li et al. 2019).

Here, we identified fixed indels that are uniformly present in all variants and attempted to determine their generation mechanisms and found a considerable number of indels generated by alteration in the number of tandem repeats, which frequently encode IDRs. Our analyses also revealed that longer internal exons tend to experience more indels and generally have a higher prevalence of tandem repeats, suggesting that long internal exons were generated chiefly by repeat expansions. We propose that the expansion of tandem repeats encoding IDRs was a crucial evolutionary mechanism by which the LECA had arisen from the FECA.

## Results

### Selection of fixed indels in internal exons

Exploiting a wealth of sequences in the Ensembl database (Martin et al. 2023), we first aligned all variants of orthologous genes in closely related sp. 1 and sp. 2 and selected sections missing in sp. 2 as indel candidates (Fig. 1A,C). We then carried out BLASTN alignments (Altschul et al. 1990) of all variants of orthologous genes in spp. 1 and 2 and an outgroup to classify them into insertion and deletion candidates; those present in an outgroup variant were considered as insertion candidates because the segments were probably absent before the bifurcation of sp. 1 and sp. 2 (Fig. 1B), while those missing in an outgroup variant were regarded as deletion candidates as the segments were likely to be present in the immediate ancestor of sp. 1 and sp. 2 (Fig. 1D). To filter out candidates that correspond to splicing variants, we applied uniformity tests; we selected the insertion candidates whose segments are present in all sp. 1 variants and are unexceptionally absent in sp. 2 and outgroup variants and regarded them as fixed insertions (Fig. 1B); we likewise chose the deletion candidates whose segments are invariably absent in sp. 2, but omnipresent in both sp. 1 and outgroup variants and considered them as fixed deletions (Fig. 1D). Finally, only those in internal exons are selected and are called fixed indels. We present the actual steps followed to select fixed indels in internal exons and the number of cases in Supplemental Fig. S1 and Table S1. (For explanations of step 5, see below.)

**Figure 1.**
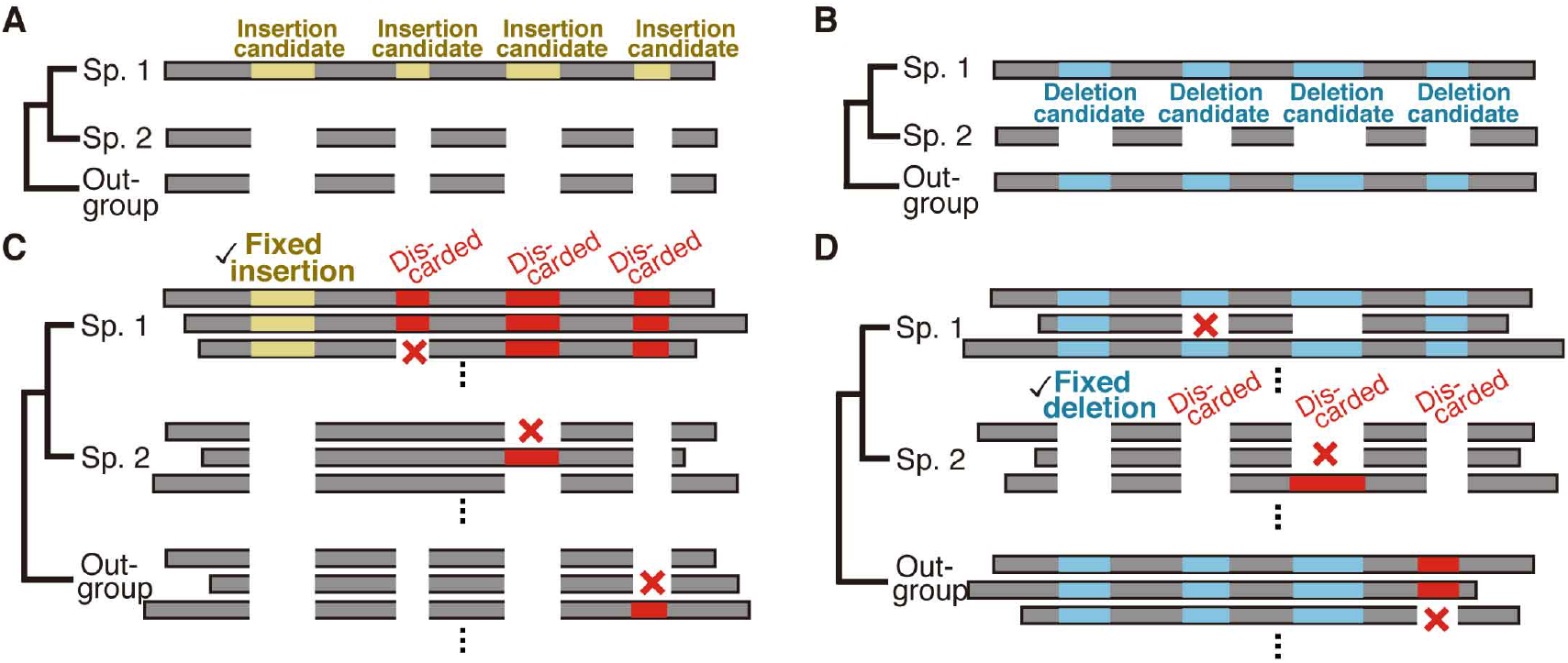
Selection of fixed indels. Rectangles represent sections of coding sequences and are arranged so that aligned segments match vertically. (*A, B*) The criteria for selecting fixed insertions and (*C,D*) fixed deletions are schematically shown.

Insertions are probably generated either by intron to exon conversion (exonization) (Fig. 2A-H) or by genomic modifications, the latter of which may be caused by expansion of tandem repeats (Fig. 2i), homologous recombination (Fig. 2J), or insertion of exogenous segments (Fig. 2K). Inserted segments that are presumably generated by exonization but correspond to introns with lengths less than 70 nt (Fig. 2B,F) were discarded (step 5 in Supplemental Fig. S1) because such introns may be imperfectly removed (Abebrese et al. 2017), making them unqualified as “fixed” insertions. Similarly, the three generating mechanisms of deletions are exon to intron conversion (intronization) (Fig. 2I-S), contraction of tandem repeats (Fig. 2T), homologous recombination (Fig. 2U), and genomic deletion (Fig. 2V). Deleted segments that corresponded to short (<70 nt) introns (Fig. 2M,Q) were removed for the aforementioned reason. (In the following, we additionally carried out analyses of the permissive sets of fixed indels to demonstrate that the removal does not affect conclusions.) Insertions with corresponding introns in sp. 2 possessing no significant homology were considered as possibly generated by exonization (Fig. 2A,C,D), while those alignable to intron segments in sp. 2 were classified as probably generated by exonization (Fig. 2E,G,H). As introns generally have a higher mutation rate than exons do (Li 1997), exons created by exonization may no longer possess detectable sequence homology with the introns. Likewise, we consider deletions as possibly generated by intronization if the corresponding intron segment in sp. 2 has no discernable similarity to the exons in sp. 1 (Fig. 2L,N,O), while sorting those as probably generated by intronization if the intron segments were alignable to the exons (Fig. 2P,R,S). We distinguish fixed insertions of entire exon(s) (Fig. 2C,G) from those within exons (Fig. 2A,C) and regard fixed deletions of entire exon(s) (Fig. 2N,R) separately from those within exons (Fig. 2L,P). The total number of indels within internal exons as well as their breakdown by indel location within exons are shown in Supplemental Tables S2 and S3, the latter of which list numbers without removing those corresponding to short (<70 nt) introns (designated the “permissive” sets of fixed indels). Fixed indels of entire exon(s) are rare and almost all the fixed indels in internal exons are in the middle of rather than at 5’ and 3’ ends of exons (Supplemental Fig. S2).

**Figure 2.**
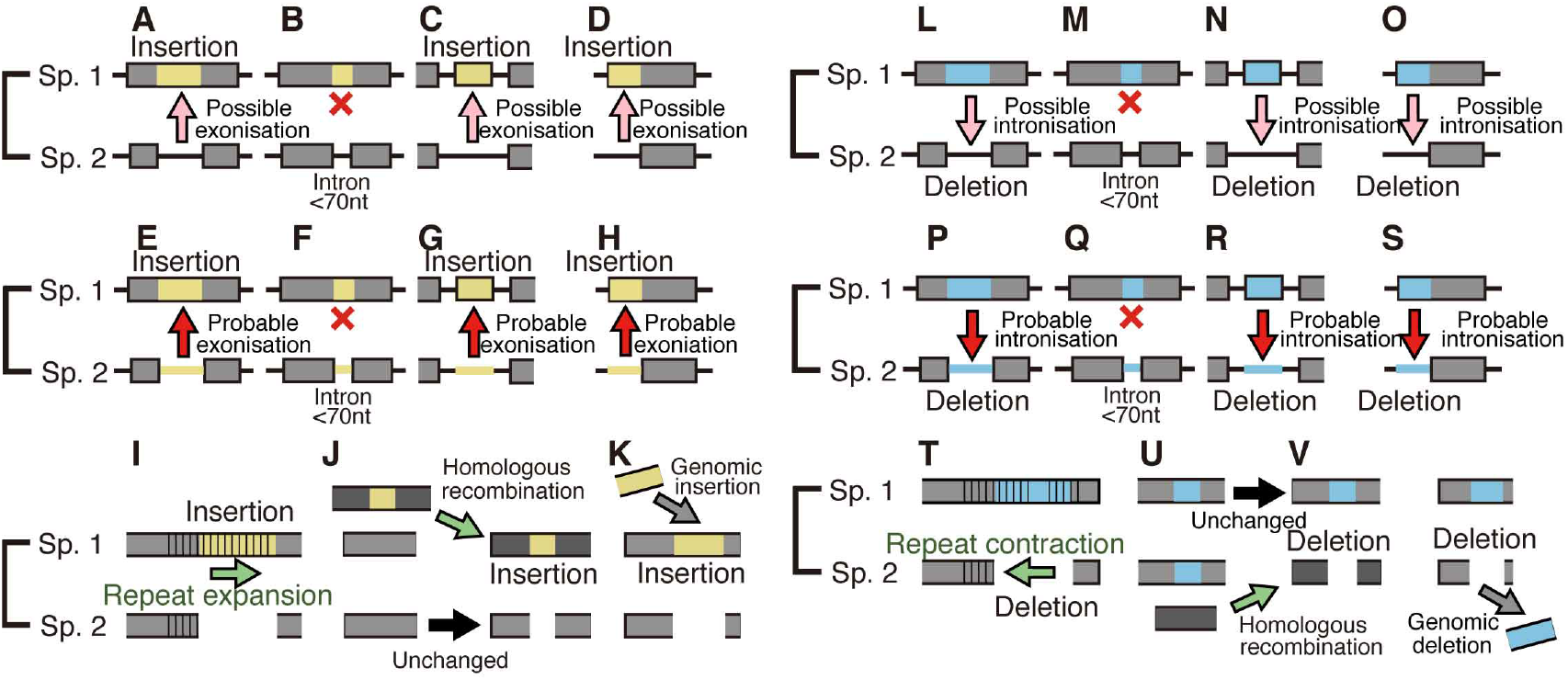
Generation mechanisms of fixed indels. Presumed generating mechanisms of fixed indels are pictorially explained. Gray rectangles and horizontal bars stand for exons and introns, respectively, while parallel vertical bars signify tandem repeats.

Six examples of fixed indels are provided (Fig. 3) and two examples of indels that were removed due to their correspondence to excessively short (<70 nt) introns are shown (Supplemental Fig. S3). The human protein in Fig. 3A is a protein enabled homologue involved in a range of processes dependent on cytoskeleton remodeling and cell polarity, the *D. melanogaster* protein in Fig. 3B is E2F transcription factor, isoform D, whereas the rat protein presented as Fig. 3C is nuclear factor erythroid 2-related factor 2, which is a transcription factor that plays a key role in the response to oxidative stress. The chimpanzee protein depicted in Fig. 3D is a trinucleotide repeat-containing protein that plays a role in RNA-mediated gene silencing, the human protein shown in Fig. 3E is GRIP and coiled-coil domain-containing protein isoform 2, while the human protein in Fig. 3F is SMG1 phosphatidylinositol 3-kinase-related kinase partaking in both mRNA surveillance and genotoxic stress response pathways.

**Figure 3.**
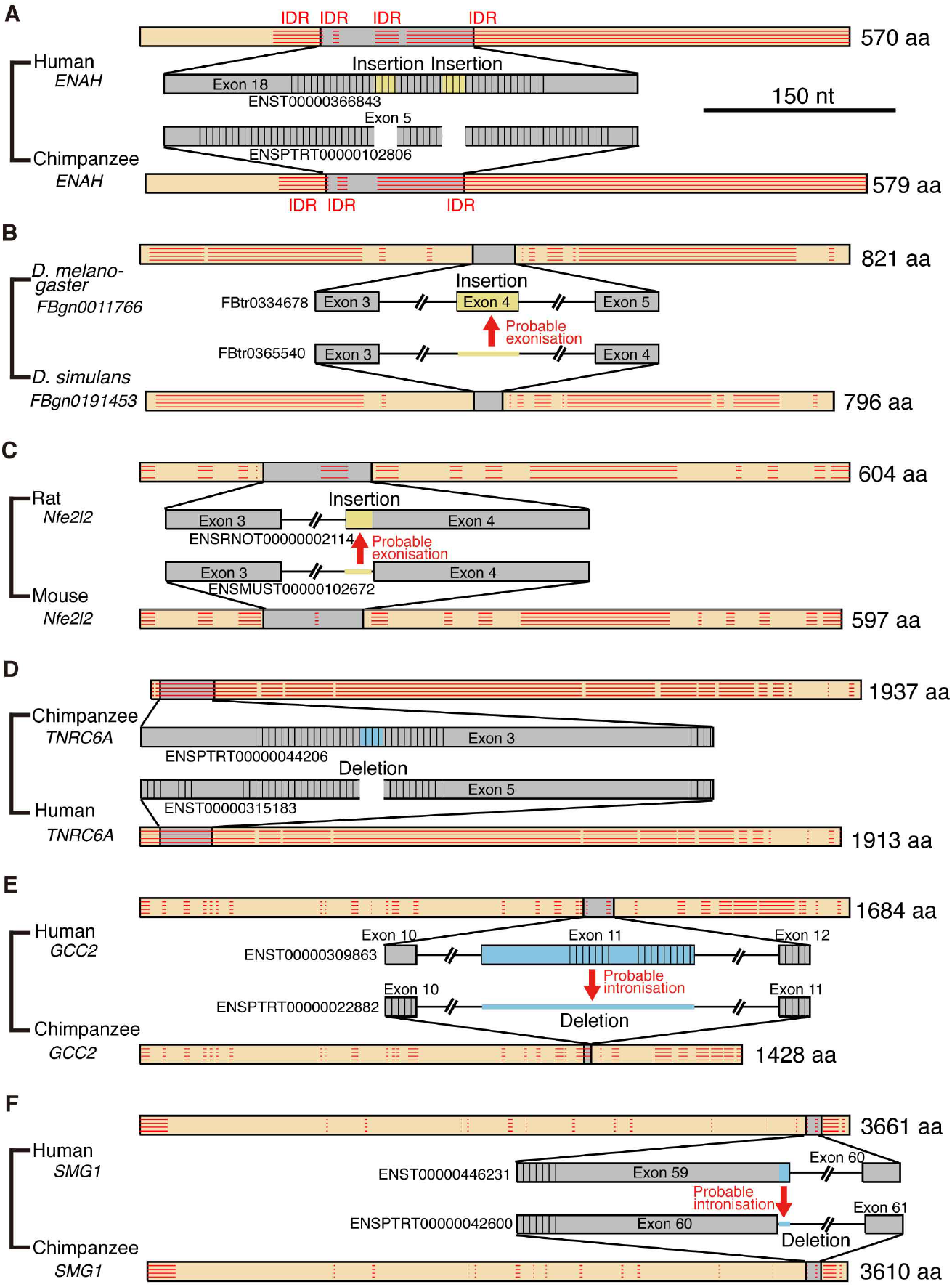
Actual examples of fixed indels. Examples of fixed insertions (*A*-*C*) and deletions (*D*-*F*) depicted as in Fig. 2 in the scale indicated with tandem repeats represented by parallel vertical bars. The gene IDs are shown in italics, while the encoded proteins are drawn as orange rectangles in arbitrary scales with horizontal parallel bars representing IDRs and gray rectangles corresponding to the depicted genomic sections..

### Generation mechanisms of fixed indels

The residue-wise distributions of generation mechanisms of fixed indels are graphically presented (Fig. 4) with the indels whose generating mechanisms remain unidentified labeled “unknown”. No indel cases involving transposable elements were detected even if the expectation value of the BLASTN search was relaxed to 10 and no insertion cases whose generation mechanisms are unknown were long enough to be assessed for involvement of other exogenous elements. The corresponding figure of the permissive sets of fixed indels is presented as Supplemental Fig. S4. The numbers of fixed indel events and those of residues in insertion and deletion are tabulated (Supplemental Table S4), while the corresponding numbers of the permissive sets of indels are presented as Supplemental Table S5. Although interspecies variations are considerable, on average 51% of inserted and 40% of deleted residues were attributable to alterations in tandem repeats and the generation mechanisms of a substantial number of fixed indels remain unidentified (see Discussion). Fixed indels generated by intronization and exonization exist but have small shares except in human insertions, chimpanzee deletions, and rat indels, whereas those attributable to homologous recombination are universally rare.

**Figure 4.**
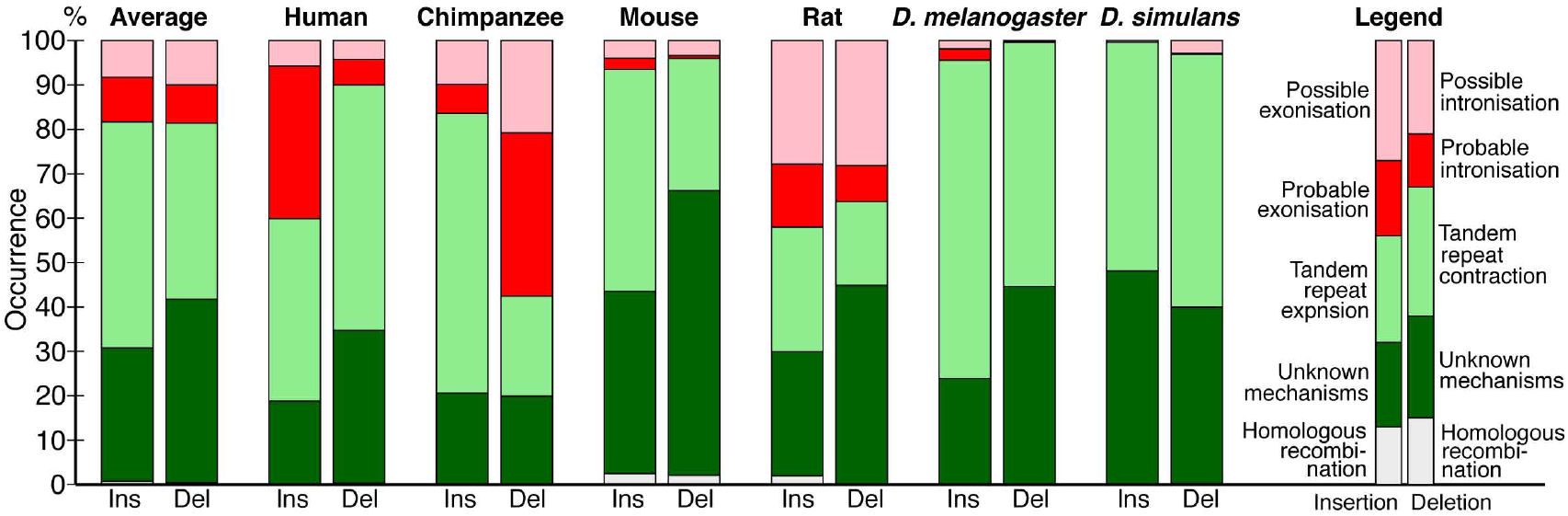
Generating mechanisms of fixed indels.

### Long internal exons tend to have a high incidence of indels

As we previously found that long internal exons tend to encode IDRs (Fukuchi et al. 2023), we sought to search for generation mechanisms of long internal exons and thus determined the dependence of indels on exon length. We first calculated the abundance of internal exons in 30 nt length bins (Supplemental Fig. S5) and found that, while long internal exons are rare in the four mammals, the two *Drosophila* species showed length distributions shifted rightward, confirming a previous report (Hawkins 1988). We then examined the frequencies of indels in each length bin of internal exons, using the length of the corresponding sp. 1 exons for the analyses of insertion in sp. 1 and deletion in sp. 2. The length of the internal exon with an insertion utilized was that of the exon minus the insertion length because that is the presumed exon length before insertion. In the example presented as Fig. 3C in which a 21 nt insertion exists in rat exon 4 of 189 nt, we used168 nt as the length of the primordial exon in which the insertion occurred. (Although 168 nt happens to be the length of the corresponding sp. 2 exon in this case, the lengths sometimes disagree as some sp. 2 exons have been altered). The indels of entire exon(s) (Fig. 2C,G,N,R), which constitute small fractions (Supplemental Fig. S2, Tables S2 and S3), were not included in this and subsequent analyses because our aim was to analyze incremental length changes in internal exons. Interestingly, we found higher frequencies of indels in long internal exons without exception (Fig. 5 and Supplemental Fig. S6). The correlation coefficients significantly differ from 0 (*P*-value<10^−2^) for insertions in *Drosophila melanogaster* and all deletions except for those in human and chimpanzee. Also, deletion frequency tends to be higher than insertion frequency in each species except in short intron length bins of human insertions. That deletion generally exceeds insertion agrees with previous reports (de Jong and Rydén 1981; Redelings et al. 2024).

**Figure 5.**
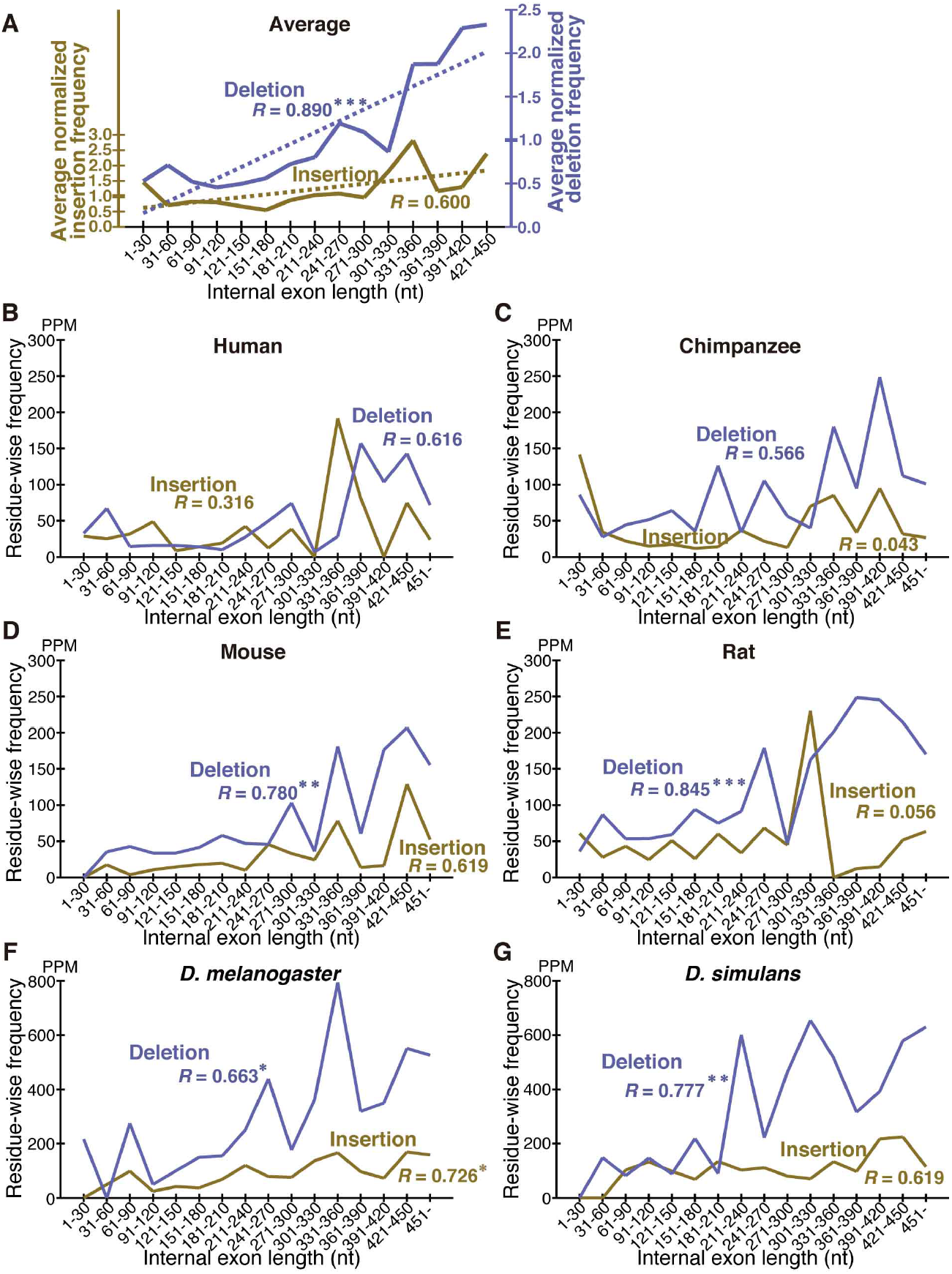
Long internal exons tend to have a high incidence of indels in each species. One, two, and three asterisks signify that the correlation coefficient is significantly different from zero at *P*-value<10^−2^, 10^−3^, and 10^−4^, respectively. (*A*) The normalized averages of insertion (left scale) and deletion (right scale) frequencies are shown with the scale of deletion frequencies multiplied by the average ratio of deletion to insertion frequency, which is the arithmetic average of the ratios of deletion to insertion frequency of the six species. The dotted lines are regression lines for insertion (yellow) and deletion (blue) frequencies. (*B*-*G*) The indel frequencies in each internal length bin in each species are graphed.

As the smallness of indel residues seemed to cause fluctuations in indel frequency, we calculated the average indel frequency of the six species in each internal exon length bin. The combined dependence of indel frequency on internal exon length (Fig. 5A and Supplemental Fig. S6A) shows that positive, statistically significant correlations exist, and that deletion frequency is on average higher than insertion frequency with the difference more pronounced in long internal exons. The latter observation reflects the fact that insertion frequency mostly levels off in the long length range, while deletion frequency does not.

### Most fixed indels encode intrinsically disordered regions, conserve reading frames, and are short

As indels reportedly occur preferentially in IDRs (Light et al. 2013A, B; Khan et al. 2015), we calculated the fractions of IDRs in the fixed indels. For deleted segments in sp. 2, we used the corresponding segments in sp. 1 (Fig. 1D) as a proxy. In agreement with the literature, most indels encode IDRs and the fractions are all much higher than those of all coding exons (Fig. 6A and Supplemental Figs. S7 and S8). We also examined indel lengths and verified reports (Wetterbom et al. 2006, Mills et al. 2006; Khan et al. 2015) that those of nearly all indels in coding sequences are multiples of 3 nt (Fig. 6B and Supplemental Figs. S9 and S10). Frame preserving insertions except for those containing stop codons merely add residues to the encoding proteins, while frame-preserving deletions excluding those containing start or stop codons simply remove residues. Since most indels encode IDRs, alterations in the number of amino acids caused by fixed indels are likely to be tolerated. The length distributions (Fig. 6C and Supplemental Figs. S11 and S12) demonstrate that shorter indels predominate, in agreement with reported findings (Benner et al. 1993; Qian and Goldstein 2001; Khan et al. 2015) and almost all fixed indels are shorter than 19 nt.

**Figure 6.**
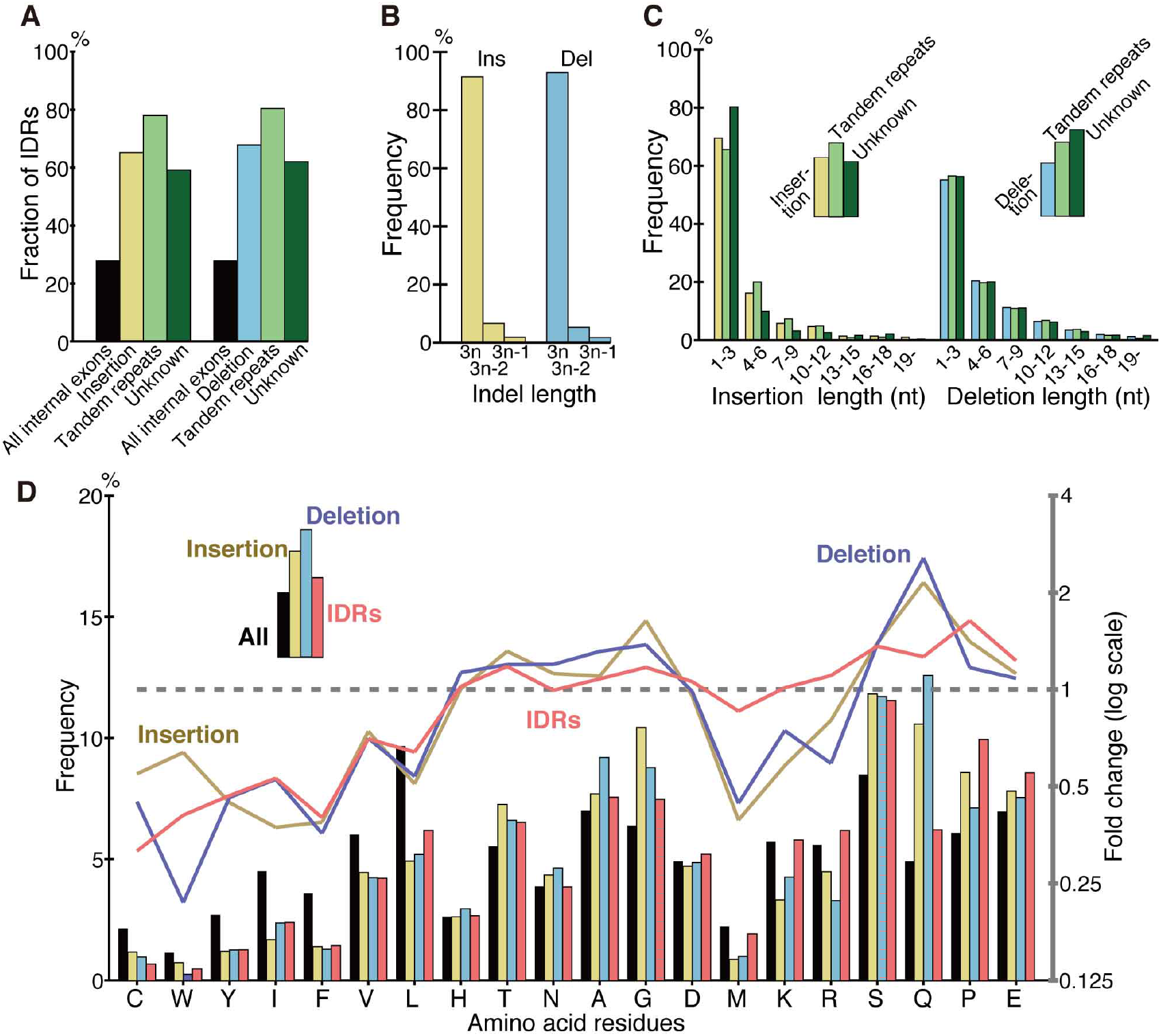
Most indels encode IDRs, are multiples of 3 nt in length, and are short, and amino acid compositions of indels resemble those of IDRs. The bars and line graphs represent arithmetic averages of the six species. (*A*-*C*) The label unknown signifies that the generation mechanisms are unknown. (*D*) The frequency of each amino acid encoded by all internal exons (“All”), that of fixed indels, and that of IDRs are represented by bars (left scale). The line graphs (right scale) are the fold changes in frequency relative to that of all internal exons.

### Amino acid compositions of indels resemble those of IDRs

As most indels encode IDRs and IDRs have a characteristic amino acid composition (Dunker et al. 2008), we calculated the amino acid compositions of indels (Fig. 6D, Supplemental Fig. S12). Indels have amino acid compositions similar to those of IDRs in that order-promoting residues are depleted, while disorder-promoting residues appear more frequently than the average. For the data presented, the correlation coefficient between log fold changes of insertion and those of IDR is 0.776 (statistically significant at *P*-value<10^−4^), while the correlation coefficient between log fold changes of deletion and those of IDR is 0.826 (statistically significant at *P*-value<10^−5^).

### Long internal exons tend to have tandem repeats

What accounts for the higher incidences of indels in longer internal exons? We found no grounds to suppose that intronization, exonization, and homologous recombination occur more frequently in longer internal exons. Since tandem repeats account for a large fraction of indels (Fig. 4) and longer internal exons generally experience more indels (Fig. 5), we thought it possible that tandem repeats are more prevalent in long internal exons. We thus calculated the fraction of each internal exon occupied by tandem repeats and investigated its dependence on internal exon length. The results (Fig. 7) verified the conjecture; the longer internal exons are, the higher the repeat frequency with all the correlation coefficients statistically significant at *P*-value<10^−2^ and the conclusion remains unchanged in the longer length range (1-840 nt instead of 1-450 nt) (Supplemental Fig. S14).

**Figure 7.**
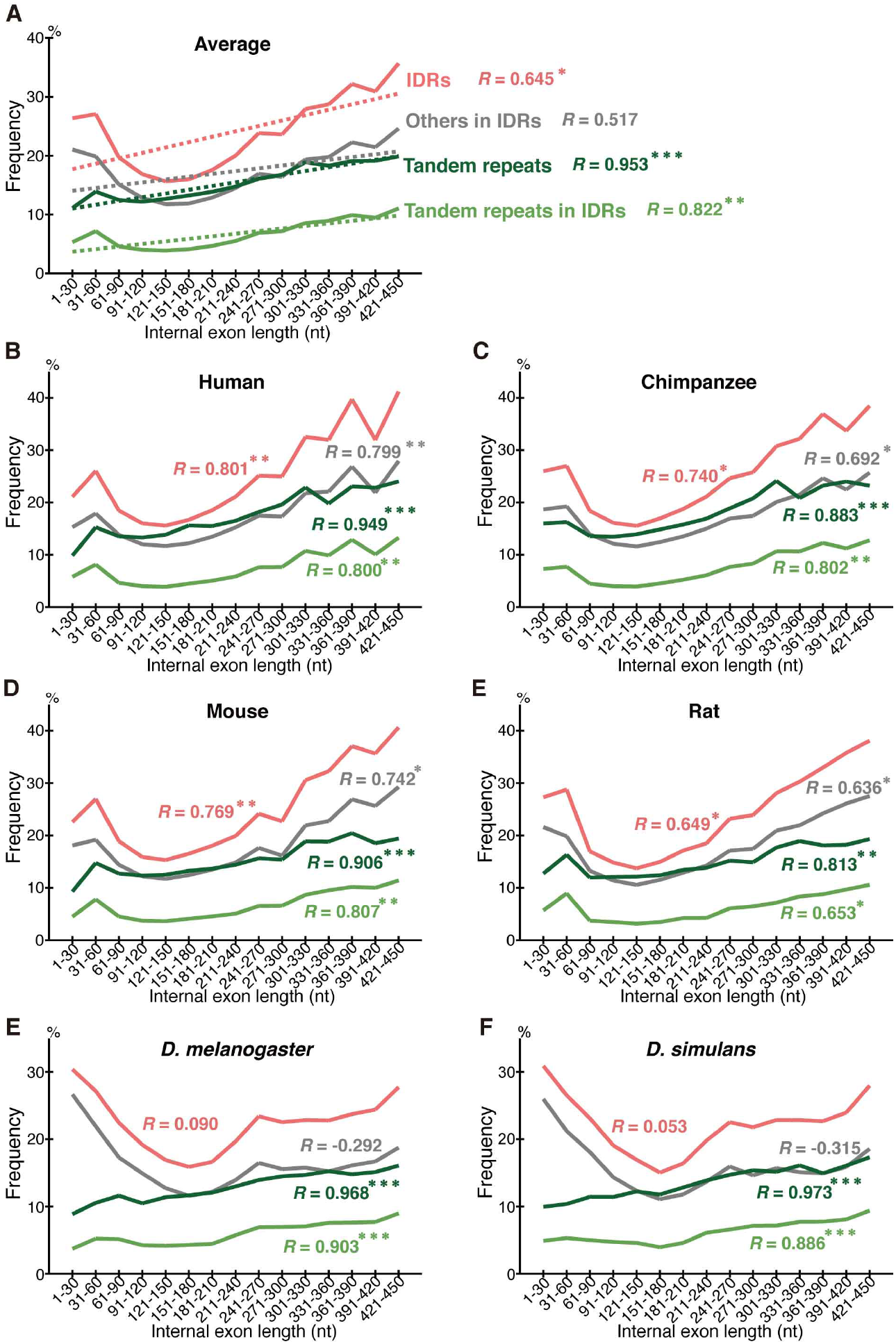
Long internal exons tend to have tandem repeats as well as IDRs. The fractions of tandem repeats, IDRs, tandem repeats in IDRs, and others in IDRs are shown with the correlation coefficients of the fractions with exon length. Asterisks signify statistical significances as in Fig. 5 legend. (*A*) Arithmetic averages are plotted with regression lines (dotted lines).

To examine how much of the higher prevalence of IDRs in longer internal exons is explained by tandem repeats, we calculated the fractions of IDRs, tandem repeats within IDRs, and others within IDRs (i.e., IDRs that are not in tandem repeat segments) in each length bin in the six species in the 1-450 nt (Fig. 7) and 1-840 nt length ranges (Supplemental Fig. S14). Note that the previously reported dependence of the fraction of IDRs on internal exon length (Fukuchi et al. 2023) was more pronounced in the longer exon length range. Since the IDRs are divided into tandem repeats (within IDRs) and others, the sum of the fractions of the two constituents is equal to that of IDRs and, consequently, the sum of the slopes of the regression lines of the two is identical to that of IDRs. If the positive correlation of IDRs are entirely accounted for by tandem repeats, the slope of the regression line of tandem repeats within IDRs will be the same as that of IDRs, whereas the slope of others will be zero, that is, the fraction of others will be constant irrespective of intron length. The observed slope of the regression line of the fraction of tandem repeats on IDRs was 0.438 per cent/unit, where 1 unit equals 30 nt, and that of others in IDRs was 0.478, while that of the corresponding slope of the fraction of IDRs was 0.916. Thus, only 48% of the dependence of IDRs on internal exon length is contributed by tandem repeats. As tandem repeats constitute a mere 28.3% of IDRs on average, however, they make a disproportionate contribution to IDRs. On the other hand, in the longer range of exon length (Supplemental Fig. S14A), the contribution ratio of tandem repeats in IDRs was 36%, lower than 48%.

### The frequency of indels correlates with that of tandem repeats

Since both the frequency of indel and that of tandem repeats show positive correlation with internal exon length, the two frequencies are likely to be correlated. The correlations between the normalized insertion/deletion frequency and the normalized repeat frequency are shown by scatter plots (Fig. 8A, Supplemental Fig. S15). The correlation coefficient between insertion and repeat frequencies is 0.597 (statistically significant at *P*-value<0.05), while that between deletion and repeat frequencies is 0.885 (statistically significant at *P*-value<10^−5^). In confirmation of the expectation, the indel frequencies are significantly correlated with repeat frequency, with the deletion frequency more strongly so.

**Figure 8.**
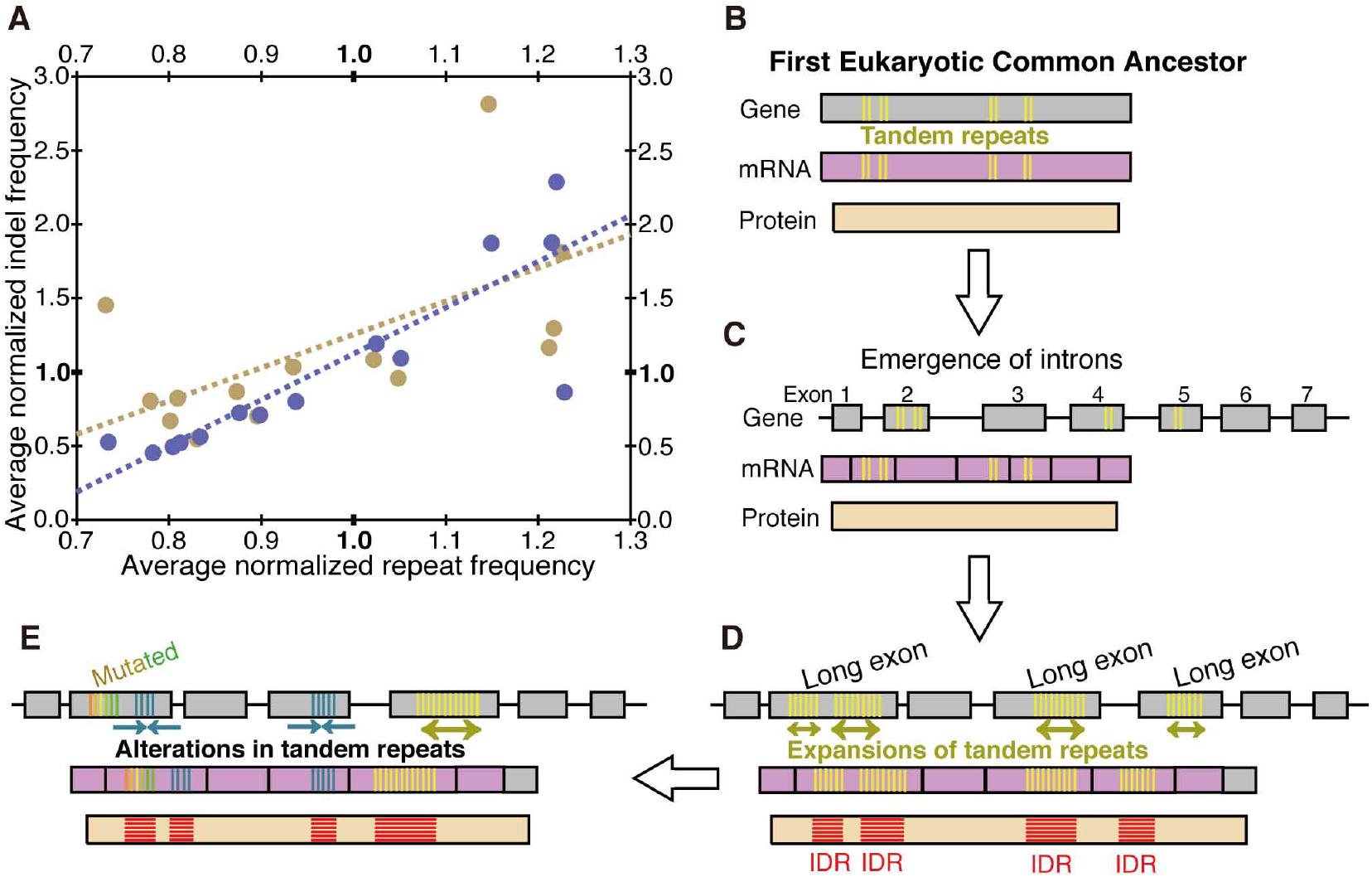
Indel frequencies are correlated with tandem repeat frequency and a proposed model. (*A*) Scatter plots of the normalized repeat frequency of each length bin and the corresponding normalized insertion (yellow dots) and deletion (blue dots) frequencies are shown. Dotted lines (yellow: insertion, blue: deletion) represent regression lines. (*B*-*D*) Proposed evolutionary steps that had led to the last eukaryotic common ancestor that have long internal exons that frequently encode IDRs. (*E*) An ongoing process.

## Discussion

We unexpectedly discovered that fixed indels occur preferentially in longer internal exons and large fractions of indels were generated by alterations in tandem repeats. We also found a high prevalence of tandem repeats in long internal exons, which accounts for the previously observed tendency of long internal exons to encode IDRs (Fukuchi et al. 2022) as tandem repeats mostly encode IDRs (Simon and Hancock 2009). In view of the findings of this research, we propose that long internal exons had resulted from primordial short exons (Fig. 8C) that had evolved from the first eukaryotic common ancestor (FECA) (Fig. 8B) mainly by expansion of tandem repeats that added IDRs to the encoded proteins, which constitutes a step toward the emergence of the last eukaryotic common ancestor (LECA) (Fig. 8D). This model explains how eukaryotic proteins acquired IDRs, especially in sections encoded by long internal exons, while only a small fraction of prokaryotic proteins consists of IDRs. This step also accounts for the increased frequency of tandem repeats in eukaryotic proteins compared to prokaryotic proteins. Moreover, we suggest that, after the LECA, long internal exons have been subject to frequent contractions and, to a lesser extent, expansions due to alterations in tandem repeats (Fig. 8E). As nearly all fixed indels are nonframeshifting (Fig. 6B), indels generated by tandem repeats mostly result in tandem repeats in proteins. Since protein tandem repeats play crucial roles in ligand binding and transcriptional regulation (Kobe and Kajava 2001; Yagi et al. 2014), indels generated by tandem repeats may have important biological significance.

The number of human indels in ALL exons identified in this investigation is the sum of the number of human insertions and that of chimpanzee deletions at step 5, and is 863 (Supplemental Table S2), which is larger than the reported number, 262, of human indels in all coding exons (Mills et al. 2006). The increase in number is attributable not only to revised sequence data we had the good fortune to use, but also to the methodology that identifies changed alternative splicing in addition to genomic alterations in constitutively spliced exons.

Although the frequencies of the detected fixed indels are small (less than 0.06%), they may well be an underestimate as our methods of selecting fixed indels probably capture only a fraction of actual fixed indels. This is both because transcripts with corresponding segments must be present in all three species in our identification method and because BLASTN alignments can be made only for highly conserved nucleotide sequences. Additionally, the outgroup requirement means that the capture rate of fixed indels varies from one species group from another and thus makes it inappropriate to compare indel frequencies of different groups; we for instance cannot meaningfully compare the indel frequencies of human and mouse. Moreover, the availability of more variant data may disqualify some fixed indels due to the presence of counterexamples in the newly identified variants. Since the methodology has no bias on exon length, however, the finding that higher indel frequency is observed in longer internal exons is unaffected by these shortcomings.

We consider the fractions of fixed indels generated by tandem repeats in exons represent an underestimate; slight sequence changes after repeat expansion and contraction make tandem repeats imperfect, making them undetectable by the repeat detection program. It is thus conceivable that indels generated by unknown mechanisms contain those attributable to tandem repeats, although they may well include some cases generated by inverted and mirror repeats (Burssed et al. 2022), which are undetectable by the program we used. In fact, the fractions of IDRs in indels generated by alterations in tandem repeats are comparable to those produced by unknown mechanisms (Fig. 6A), their length distributions are nearly identical (Fig. 6C), and amino acid compositions of the two groups are similar (Supplemental Fig. S16). It is thus plausible that many indels whose generation mechanisms remain unidentified were really generated by alterations in tandem repeats. An underestimation of tandem repeats can explain why the tandem repeats explain only 48% (in the internal exon length range 1-450 nt, Fig. 7A) or even less in the longer (1-840 nt) exon length range of the dependence of IDRs on the length of internal exons; unidentified tandem repeats may account for some of the remaining dependence.

Notwithstanding all the emphasis on tandem repeats, we would like to reiterate that intronization and exonization also generate deletions and insertions, respectively, and they account for considerable fractions of fixed insertions in *Homo sapiens*, fixed deletions in chimpanzee, and fixed indels in rat (Fig. 4). The fact that many indels were “probably’” generated by intronization and exonization makes it quite likely that they really account for some indels. The wide variation in the fractions of intronization and exonization is largely explainable by disparity in location distribution of indels within internal exons; most insertions at 5’ and 3’ ends of exons and those involving entire exons were judged to be generated by exonization (70.1, 77.2 and 58.6%, respectively), and nearly all deletions at 5’ and 3’ ends of exons and those of entire exon(s) were attributable to exonization (85.0, 91.1 and 97.9%, respectively). By contrast, intronization and exonization were seldom assigned as the generation mechanisms of indels within exons (0.7 and 1.0%, respectively). The insertions in human and rat and the deletions in chimpanzee and rat have high total proportions of those at the ends of exon and those involving entire exons (Supplemental Fig. S2), and these have comparatively high fractions of cases generated by intronization/exonization. Indel analyses of other species are needed to judge whether intronization and exonization generally generate less indels than tandem repeats do.

What accounts for the general preponderance of deletion rates over insertion rates? We consider it possible that insertion is evolutionarily disfavored especially in long internal exons because splicing of many excessively long exons (>300 nt) is inhibited (Robberson et al. 1990; Chen and Chasin 1994; Sterner et al. 1996). This interpretation dovetails with the observed tendencies that insertion frequency of internal exons initially rises but levels off at around 300 nt, while deletion frequency shows a monotonically increasing trend (Fig. 5A). However, we excluded fixed insertions that create entire exons (Fig. 2C,G) and fixed deletions that eliminate whole exons (Fig. 2N,R) from our analyses of exon length dependence and our analysis cannot capture cases of exon fusion and fission that do not affect coding sequences. Although analysis of exon evolution including these neglected cases is expected to elucidate exon evolution dynamics in its entirety, it is beyond the scope of the current study.

The existence of a positive correlation between tandem repeat frequency and internal exon length was an intriguing finding. Since four other eukaryotes, *Oryza sativa, Arabidopsis thaliana, Caenorhabditis elegans*, and *Schizosaccharomyces pombe* were shown to have a positive correlation of the fraction of IDRs and internal exon length (Fukuchi et al. 2023), we checked if repeat frequency and internal exon length are correlated in the 1-450 nt range in these model eukaryotes, too. We found the first three model eukaryotes have statistically significant (*P*-value<10^−2^) positive correlations, but *S. pombe* does not (Supplemental Fig. S17). The case of *S. pombe* is apparently inconsistent with the notion that long internal exons were produced mainly by the expansion of tandem repeats. However, as stated above, the fraction of tandem repeats is likely to be underestimated, and the underestimation can explain the lack of correlation in the yeast species.

As far as we are aware, this represents the first systematic identification of fixed indels including those generated by constitutive alterations in alternative splicing and examination of the dependence of their frequencies on internal exon length. As our observations are limited to indels in six species only, however, the generality of our findings needs to be tested by indels in other species. Unfortunately, it is not easy to apply the current methodology to many other species since the current selection method requires the presence of a trio of closely related species with completely sequenced genomes and an abundance of sequenced variants. For instance, the application of our methods to *C. elegans, C. briggsae* with *C. japonica* as the outgroup identified less than 70 cases of fixed insertions in *C. elegans*, a number judged too small for further analyses. Hopefully, widespread availability of variant and genome sequences will make extensive testing possible.

## Methods

### Data sets

For the selection of *Homo sapiens* and *Pan troglodytes* (chimpanzee) indels, we chose *Gorilla gorilla* as the outgroup. As the outgroup of *Mus musculus* (mouse) and *Rattus norvegicus* (rat), we used *Peromyscus maniculatus*, while *Drosophila yakuba* served as the outgroup of *Drosophila melanogaster* and *Drosophila simulans*. We downloaded all the sequence and exon data, and the lists of orthologous genes except for that between *D. melanogaster* and *D. simulans* from the Ensembl database (Martin et al. 2023). As a list of orthologous genes between the two *Drosophila* species was unavailable from the Ensemble database, we selected pairs of orthologous genes by mutual best hit of BLASTN alignments of the longest variants. The list of transposable elements used was in release 19 of the TREP Transposable Element Platform (Wicker et al. 2002).

### Selection of fixed indels and their classification by location

The actual selection steps followed are schematically shown (Supplemental Fig. S1). We performed BLASTN alignments (Altschul et al. 1990) with the default parameters except that we set the gap-opening penalty at 10 and changed the expectation value to 10^−3^, unless otherwise stated. We identified indel candidates in BLASTN alignments with special considerations for sections of multiple alignments; we regarded two aligned sections continuous if the last query residue of one section coincides with the first query residue of the other within 3 nt. (The 3 nt allowance was made to cope with the uncertainty in alignments.) We judged indel sections identical if their genomic addresses agreed within 3 nt. In the uniformity tests, we disregarded variants that do not have sections covering indels. An indel was regarded as coinciding with the 5’ or 3’ end of an exon of sp. 1 if the start or the end of the segment coincides with the start or the end of the exon within 3 nt. We judged an indel without coincidence with exon border(s) to be located inside an exon.

### Generation mechanisms of fixed indels

An insertion is considered to be “possibly” generated by exonization if an intron whose length is more than or equal to two thirds of the length of the insertion exists in sp. 2 at the corresponding location but is upgraded to “probably” generated by exonization if at least two thirds of the residues were aligned (Fig. 2A,C-E,G,H). Likewise, a deletion is regarded as “possibly” generated by intronization if the corresponding segment in sp. 2 has an intron of at least two thirds of the length of the deleted segment but is reclassified as “probably” generated by intronization if the aligned segment is more than or equal to two thirds of the length of the deleted segment (Fig. 2L,N-P,R,S).

We identified the generation mechanisms of the rest of the fixed indels as follows. Homologous recombination is chosen as the generation mechanism of an insertion in sp. 1 based on the following conjecture (Supplemental Fig. S18); sequence A with insertion generated by homologous recombination in sp. 1 is probably more like sequence A’ in sp. 1 that replaced the sequence than to the orthologous sequence B in sp. 2. Based on this idea, we set two criteria for an insertion to be a product of homologous recombination. Firstly, there must be at least one sequence A’ of a different gene in sp. 1 aligned to the sp. 2 ortholog B, and secondly the fraction of the identity between the sequences of the insertion-containing exon and A’ is not significantly lower than that of the exon and B sequences (chi-square test). Similarly, a deletion in sp. 2 is considered to have been generated by homologous recombination if 1) sequence B in sp. 2 with an exon containing the deletion has at least one sequence B’ mapped to a different gene in sp. 2 that is aligned to the sp. 1 ortholog A and 2) the fraction of sequence identity between the exon and B’ is not significantly lower than that between the exon and A (chi-square test).

Tandem repeats were identified in coding sequences by Tandem Repeats Finder (Benson 1999) with the mismatch and indel penalties and maximum period size set at 5, 3, and 2000, respectively, while the other parameters were at default settings. If there are segments of tandem repeats in the fixed indels, the tandem repeats were regarded as responsible for generating the indels.

### Assignment of IDRs and statistical analyses

We ran IUPred3 (Erdős et al. 2021) and judged amino acids with a score more than or equal to 0.502 to be in IDRs. We carried out all the tests of statistical significance by t-test unless otherwise stated. For the data shown in Figs. 5A, 8A and Supplemental Figs. S6A and S15A, we first divided the frequency of each length bin by the average frequency of the species and arithmetically averaged the normalized values of the six species.

### Data access

All data and code necessary to reproduce these analyses are available at Figshare; the data as well as a program list, the programs for the primates, rodents, *Drosophila* species, and the remaining four eukaryotes have been respectively deposited at https://figshare.com/account/articles/27366162, https://figshare.com/account/articles/27410016, https://figshare.com/account/articles/27411264, https://figshare.com/account/articles/27411279, and https://figshare.com/account/articles/27411291.

## Acknowledgments

This work was supported in part by Grant-in-Aid for Transformative Research Areas ‘Multifaceted proteins: Expanding and transformative protein world’ from the Ministry of Education, Culture, Sports, Science and Technology in Japan, 20H05932.

