## Supplemental tables and figs. for "Longer internal exons tend to have more tandem repeats and more frequently experience insertions and deletions that are mostly in intrinsically disordered regions of the encoded proteins"

### Supplemental tables and figures

#### List of supplemental tables and figures

**Supplemental Table S1.** Number of cases at each selection step.

**Supplemental Table S2.** Numbers of cases and residues of fixed indels classified by location with respect to internal exons.

**Supplemental Table S3.** Numbers of cases and residues of fixed indels of the permissive sets classified by location with respect to internal exons.

**Supplemental Table S4.** Numbers of events and residues in fixed indels classified by generating mechanism.

**Supplemental Table S5.** Numbers of events and residues in fixed indels of the permissive sets classified by generating mechanism.

**Supplemental Figure S1.** Selection steps of fixed indels.

**Supplemental Figure S2.** Most fixed indels in internal exons are located in the middle of exons.

**Supplemental Figure S3.** Actual indel cases discarded because of their correspondence to excessively short introns.

**Supplemental Figure S4.** Distribution of generating mechanisms of the permissive sets of fixed indels.

**Supplemental Figure S5.** Length distribution of all internal exons.

**Supplemental Figure S6.** Dependence of the frequency of the permissive sets of fixed indels on the length of internal exons.

**Supplemental Figure S7.** Indels mostly encode IDRs in all species.

**Supplemental Figure S8.** Fixed indels of the permissive sets mostly encode IDRs.

**Supplemental Figure S9.** Nearly all fixed indels preserve the reading frame.

**Supplemental Figure S10.** Almost all fixed indels of the permissive sets preserve the reading frame.

**Supplemental Figure S11.** Length distributions of fixed indels in each species.

**Supplemental Figure S12.** Length distributions of the permissive sets of fixed indels.

**Supplemental Figure S13.** Amino acid compositions of the permissive sets of fixed indels resemble those of IDRs.

**Supplemental Figure S14.** Long internal exons tend to have a high fraction of repeats as well as IDRs.

**Supplemental Figure S15.** Indel frequencies of the permissive sets and repeat frequency are correlated.

**Supplemental Figure S16.** Amino acid compositions of all internal exons, those generated by repeats, and those whose generation mechanisms are unknown.

**Supplemental Figure S17** Long internal exons tend to have a high fraction of tandem repeats in *O. sativa*, *A. thaliana*, and *C. elegans*, but not in *S. pombe*.

**Supplemental Figure S18.** Criteria for selecting indel cases generated by homologous recombination.

**Supplemental Table S1.** Number of cases at each selection step.

| Insertion | Human | Chimpanzee | Mouse | Rat | ***Drosophila melanogaster*** | ***Drosophila simulans*** |
|---|---|---|---|---|---|---|
| Step 1 | 36075 | 42139 | 5835 | 4837 | 42210 | 41041 |
| Step 2 | 2246 | 1440 | 2227 | 1882 | 4321 | 3153 |
| Step 3 | 912 | 356 | 367 | 463 | 1071 | 706 |
| Step 4 | 250 | 296 | 339 | 390 | 911 | 621 |
| Step 5 | 201 | 296 | 339 | 390 | 909 | 616 |
| Step 6 | 110,126 | 175,175 | 147,147 | 215,215 | 431,432 | 303,307 |
| **Deletion** | **Human** | **Chimpanzee** | **Mouse** | **Rat** | ***Drosophila melanogaster*** | ***Drosophila simulans*** |
| Step 1 | 42139 | 36075 | 4837 | 5835 | 41041 | 42210 |
| Step 2 | 1440 | 2246 | 1882 | 2227 | 3153 | 4321 |
| Step 3 | 304 | 922 | 768 | 988 | 1640 | 2228 |
| Step 4 | 295 | 842 | 757 | 973 | 1629 | 2196 |
| Step 5 | 294 | 662 | 757 | 939 | 1623 | 2183 |
| Step 6 | 170,171 | 395,455 | 357,357 | 430,454 | 796,799 | 1083,1089 |

In step 6, the number of cases obtained following all six steps is shown first, followed by the number of cases obtained skipping step 5.

**Supplemental Table S2.** Numbers of cases and residues of fixed indels classified by location with respect to internal exons.

| **Insertion case #** | **Total** | **Inside** | **5' end** | **3' end** | **Entire exon(s)** |
|---|---|---|---|---|---|
| Human | 110 | 90 | 9 | 7 | 4 |
| Chimpanzee | 177 | 142 | 10 | 23 | 2 |
| Mouse | 147 | 139 | 4 | 4 | 0 |
| Rat | 217 | 163 | 27 | 25 | 2 |
| *D. melanogaster* | 433 | 429 | 1 | 1 | 2 |
| *D. simulans* | 307 | 272 | 12 | 19 | 4 |
| **Insertion residue #** | **Total** | **Inside** | **5' end** | **3' end** | **Entire exon(s)** |
| Human | 783 | 478 | 95 | 171 | 39 |
| Chimpanzee | 671 | 466 | 38 | 125 | 42 |
| Mouse | 600 | 561 | 12 | 27 | 0 |
| Rat | 1373 | 758 | 280 | 260 | 75 |
| *D. melanogaster* | 1897 | 1810 | 3 | 3 | 81 |
| *D. simulans* | 1579 | 1272 | 118 | 129 | 60 |
| **Deletion case #** | **Total** | **Inside** | **5' end** | **3' end** | **Entire exon(s)** |
| Human | 170 | 150 | 9 | 11 | 0 |
| Chimpanzee | 395 | 331 | 32 | 21 | 11 |
| Mouse | 357 | 339 | 8 | 9 | 1 |
| Rat | 425 | 380 | 27 | 16 | 2 |
| *D. melanogaster* | 795 | 783 | 7 | 5 | 0 |
| *D. simulans* | 1080 | 1069 | 5 | 5 | 1 |
| **Deletion residue #** | **Total** | **Inside** | **5' end** | **3' end** | **Entire exon(s)** |
| Human | 891 | 769 | 60 | 62 | 0 |
| Chimpanzee | 3348 | 1352 | 274 | 297 | 1425 |
| Mouse | 1867 | 1771 | 30 | 45 | 21 |
| Rat | 3256 | 2159 | 185 | 174 | 738 |
| *D. melanogaster* | 5626 | 5554 | 33 | 39 | 0 |
| *D. simulans* | 7135 | 6969 | 39 | 126 | 1 |

**Supplemental Table S3.** Numbers of cases and residues of fixed indels of the permissive sets classified by location with respect to internal exons.

| Insertion case # | Total | Inside | 5' end | 3' end | Entire exon(s) |
|---|---|---|---|---|---|
| Human | 126 | 106 | 9 | 7 | 4 |
| Chimpanzee | 177 | 142 | 10 | 23 | 2 |
| Mouse | 147 | 139 | 4 | 4 | 0 |
| Rat | 217 | 163 | 27 | 25 | 2 |
| *D. melanogaster* | 434 | 429 | 1 | 2 | 2 |
| *D. simulans* | 311 | 274 | 12 | 21 | 4 |
| **Insertion residue #** | **Total** | **Inside** | **5' end** | **3' end** | **Entire exon(s)** |
| Human | 912 | 607 | 95 | 171 | 39 |
| Chimpanzee | 671 | 466 | 38 | 125 | 42 |
| Mouse | 600 | 561 | 12 | 27 | 0 |
| Rat | 1373 | 758 | 280 | 260 | 75 |
| *D. melanogaster* | 1901 | 1810 | 3 | 7 | 81 |
| *D. simulans* | 1612 | 1296 | 118 | 138 | 60 |
| **Deletion case #** | **Total** | **Inside** | **5' end** | **3' end** | **Entire exon(s)** |
| Human | 171 | 151 | 9 | 11 | 0 |
| Chimpanzee | 455 | 384 | 33 | 26 | 12 |
| Mouse | 357 | 339 | 8 | 9 | 1 |
| Rat | 448 | 402 | 27 | 17 | 2 |
| *D. melanogaster* | 798 | 783 | 9 | 6 | 0 |
| *D. simulans* | 1086 | 1073 | 6 | 6 | 1 |
| **Deletion residue #** | **Total** | **Inside** | **5' end** | **3' end** | **Entire exon(s)** |
| Human | 893 | 771 | 60 | 62 | 0 |
| Chimpanzee | 3868 | 1817 | 277 | 322 | 1452 |
| Mouse | 1867 | 1771 | 30 | 45 | 21 |
| Rat | 3385 | 2285 | 185 | 177 | 738 |
| *D. melanogaster* | 5698 | 5554 | 42 | 102 | 0 |
| *D. simulans* | 7177 | 6999 | 42 | 135 | 1 |

**Supplemental Table S4.** Numbers of events and residues in fixed indels classified by generating mechanism.

| **Insertion case #** | **All** | **Homologous recombination** | **Un-known** | **Tandem repeats** | **Probable exonization** | **Possible exonization** |
|---|---|---|---|---|---|---|
| Human | 110 | 0 | 31 | 60 | 13 | 6 |
| Chimpanzee | 177 | 0 | 40 | 114 | 9 | 14 |
| Mouse | 147 | 5 | 71 | 63 | 4 | 4 |
| Rat | 217 | 9 | 76 | 82 | 25 | 25 |
| *D. melanogaster* | 433 | 1 | 128 | 301 | 1 | 2 |
| *D. simulans* | 307 | 0 | 135 | 171 | 0 | 1 |
| **Insertion residue #** | | | | | | |
| Human | 783 | 0 | 147 | 322 | 269 | 45 |
| Chimpanzee | 671 | 0 | 138 | 423 | 44 | 66 |
| Mouse | 600 | 15 | 246 | 300 | 15 | 24 |
| Rat | 1373 | 27 | 384 | 385 | 195 | 382 |
| *D. melanogaster* | 1897 | 3 | 450 | 1360 | 48 | 36 |
| *D. simulans* | 1579 | 0 | 760 | 813 | 0 | 6 |
| **Deletion case #** | **All** | **Homologous recombination** | **Un-known** | **Tandem repeats** | **Probable intronization** | **Possible intronization** |
| Human | 170 | 1 | 68 | 87 | 7 | 7 |
| Chimpanzee | 395 | 0 | 130 | 208 | 33 | 24 |
| Mouse | 357 | 6 | 233 | 101 | 3 | 14 |
| Rat | 425 | 1 | 257 | 123 | 21 | 23 |
| *D. melanogaster* | 795 | 1 | 331 | 460 | 1 | 2 |
| *D. simulans* | 1080 | 1 | 435 | 627 | 2 | 15 |
| **Deletion residue #** | | | | | | |
| Human | 891 | 3 | 306 | 493 | 51 | 38 |
| Chimpanzee | 3348 | 0 | 667 | 755 | 1230 | 696 |
| Mouse | 1867 | 39 | 1197 | 556 | 12 | 63 |
| Rat | 3256 | 3 | 1457 | 615 | 265 | 916 |
| *D. melanogaster* | 5626 | 3 | 2506 | 3093 | 6 | 18 |
| *D. simulans* | 7135 | 18 | 2832 | 4061 | 21 | 203 |

**Supplemental Table S5.** Numbers of events and residues in fixed indels of the permissive sets classified by generating mechanism.

| Insertion case # | All | Homologous recombination | Un-known | Tandem repeats | Probable exonization | Possible exonization |
|---|---|---|---|---|---|---|
| Human | 126 | 0 | 31 | 60 | 25 | 10 |
| Chimpanzee | 177 | 0 | 40 | 114 | 9 | 14 |
| Mouse | 147 | 5 | 71 | 63 | 4 | 4 |
| Rat | 217 | 9 | 76 | 82 | 25 | 25 |
| *D. melanogaster* | 434 | 1 | 128 | 301 | 1 | 3 |
| *D. simulans* | 311 | 0 | 135 | 171 | 2 | 3 |
| **Insertion residue #** | | | | | | |
| Human | 912 | 0 | 147 | 322 | 374 | 69 |
| Chimpanzee | 671 | 0 | 138 | 423 | 44 | 66 |
| Mouse | 600 | 15 | 246 | 300 | 15 | 24 |
| Rat | 1373 | 27 | 384 | 385 | 195 | 382 |
| *D. melanogaster* | 1901 | 3 | 450 | 1360 | 48 | 40 |
| *D. simulans* | 1612 | 0 | 760 | 813 | 12 | 27 |
| **Deletion case #** | **All** | **Homologous recombination** | **Un-known** | **Tandem repeats** | **Probable intronization** | **Possible intronization** |
| Human | 171 | 1 | 68 | 88 | 7 | 7 |
| Chimpanzee | 455 | 0 | 130 | 209 | 82 | 34 |
| Mouse | 357 | 6 | 233 | 101 | 3 | 14 |
| Rat | 448 | 1 | 279 | 123 | 22 | 23 |
| *D. melanogaster* | 798 | 1 | 331 | 460 | 1 | 5 |
| *D. simulans* | 1086 | 1 | 435 | 627 | 4 | 19 |
| **Deletion residue #** | | | | | | |
| Human | 893 | 3 | 306 | 495 | 51 | 38 |
| Chimpanzee | 3868 | 0 | 667 | 764 | 1679 | 758 |
| Mouse | 1867 | 39 | 1197 | 556 | 12 | 63 |
| Rat | 3385 | 3 | 1547 | 615 | 304 | 916 |
| *D. melanogaster* | 5698 | 3 | 2506 | 3093 | 6 | 90 |
| *D. simulans* | 7177 | 18 | 2832 | 4061 | 25 | 241 |

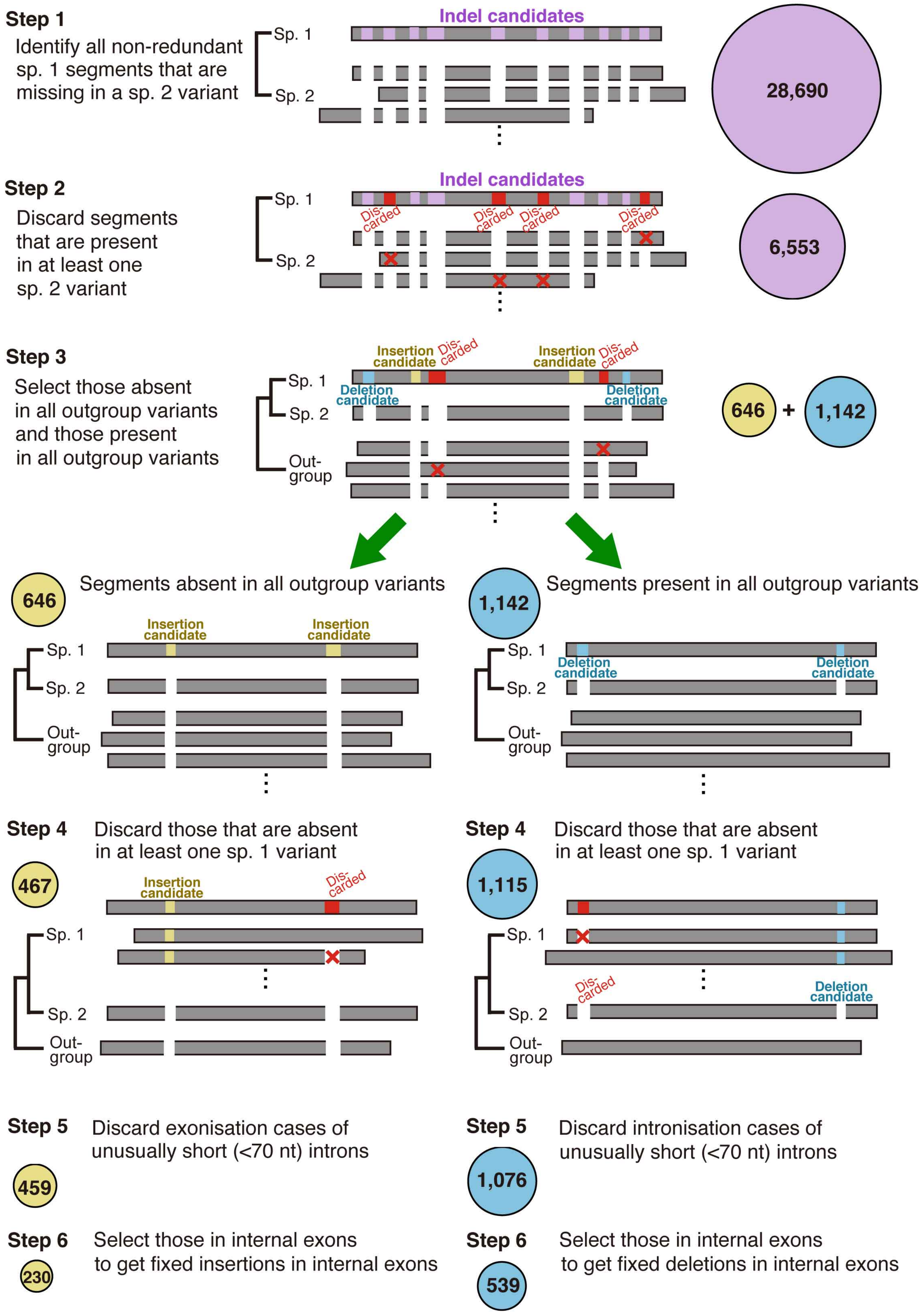


**Supplemental Figure S1.** Selection steps of fixed indels. The average numbers of cases are shown in circles.

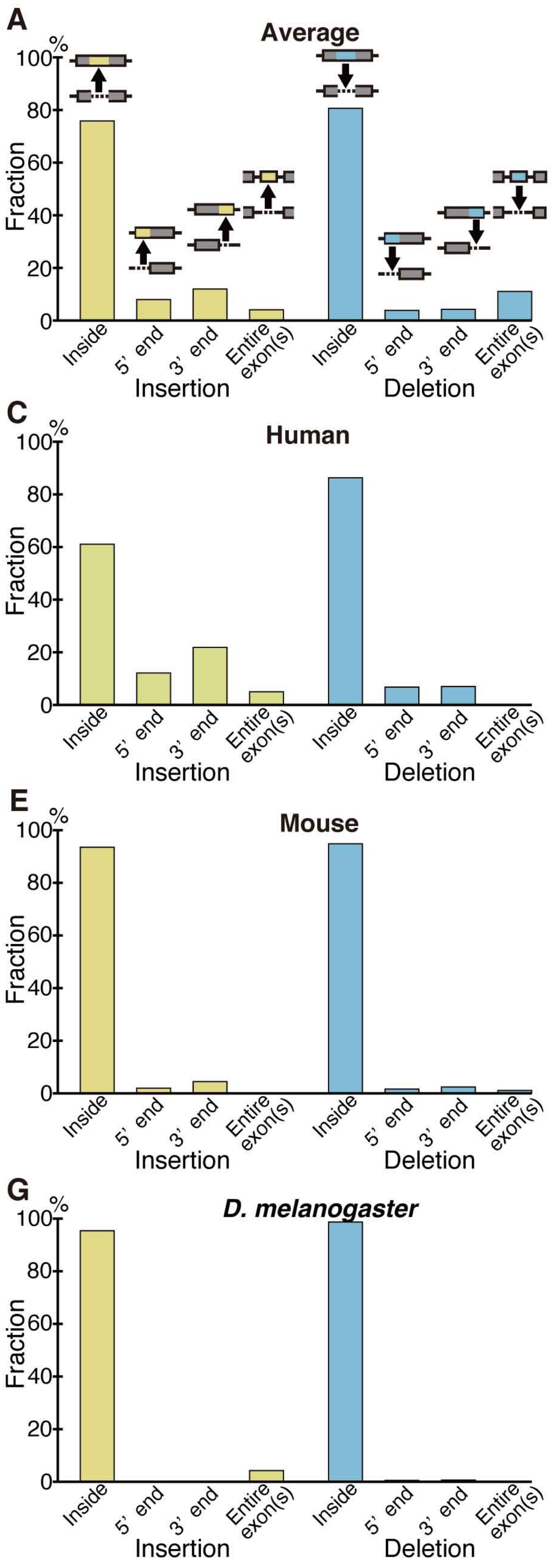


**Supplemental Figure S2.** Most fixed indels in internal exons are located in the middle of exons. (*A*) The average fractional distribution of fixed indels representing the ratio of the sum of residues in each category to the overall total. (*B*) The average fractional distribution of the permissive set of fixed indels calculated as in *A*. (*C-H*) The distribution of fixed indels of each species.

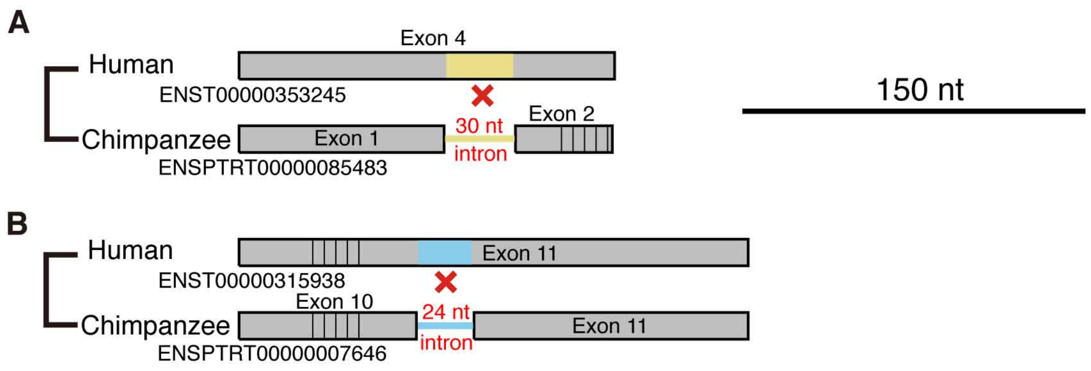


**Supplemental Figure S3.** Actual indel cases discarded because of their correspondence to excessively short introns. Legend as in Fig. 3.

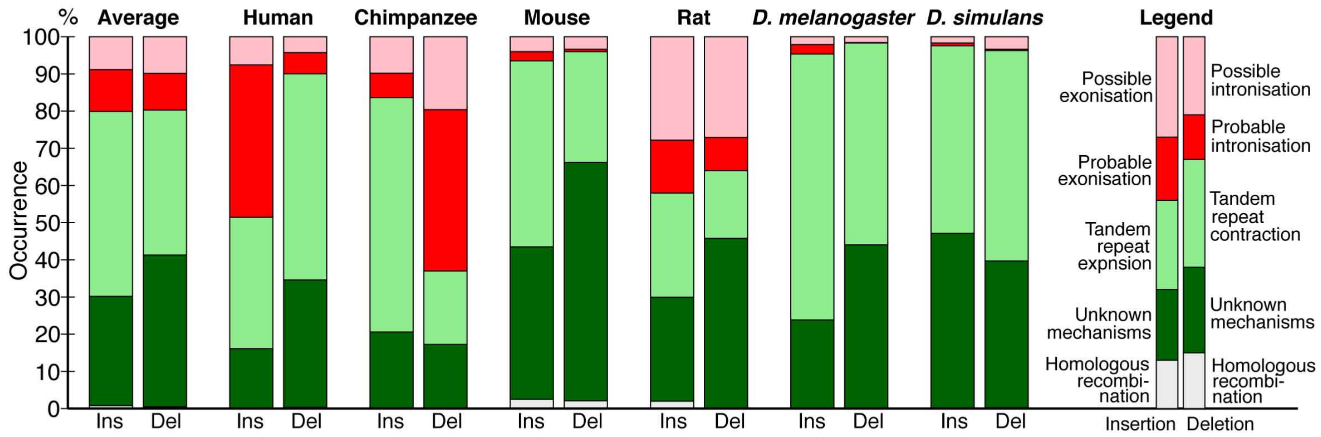


**Supplemental Figure S4.** Distribution of generating mechanisms of the permissive sets of fixed indels.

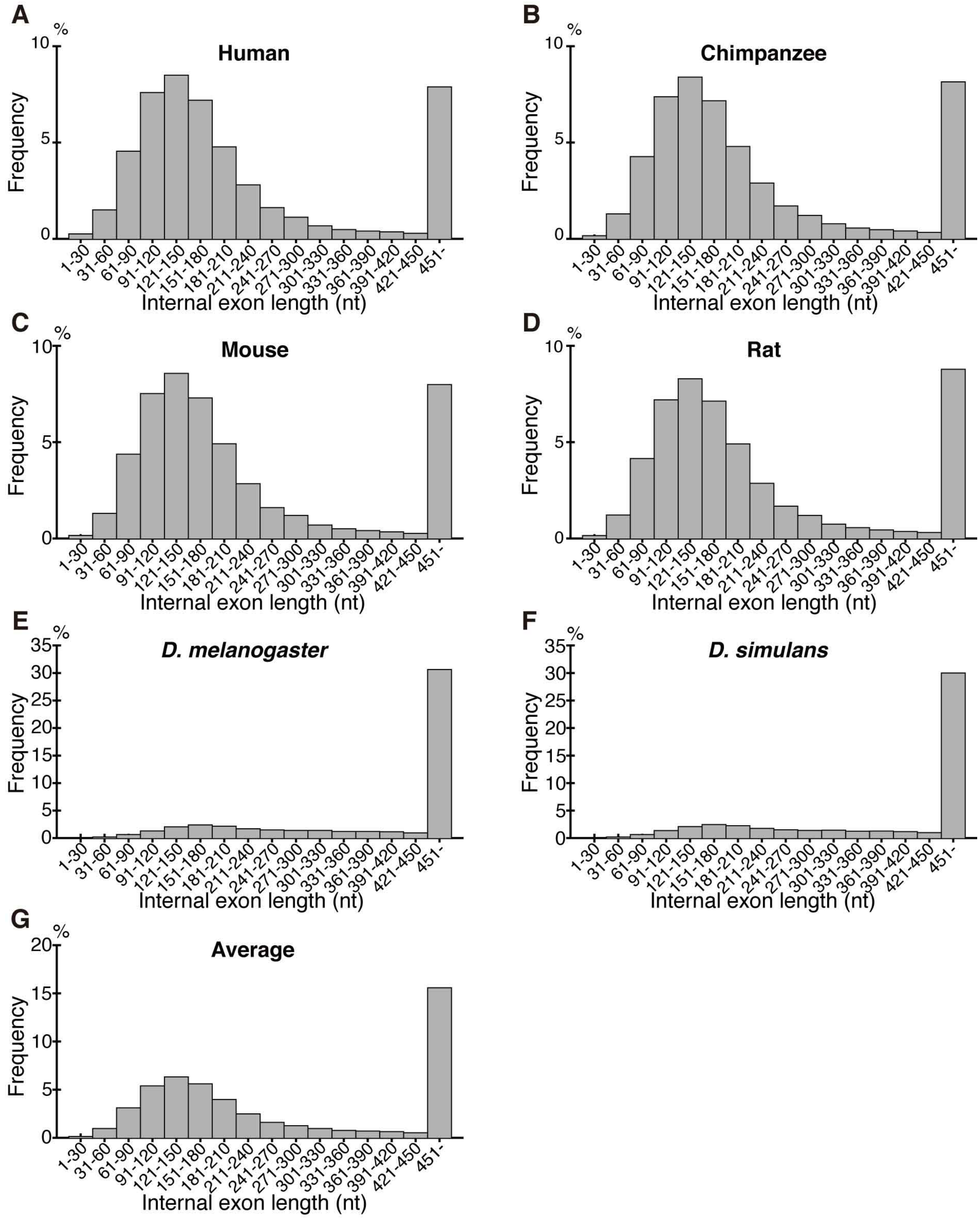


**Supplemental Figure S5.** Length distribution of all internal exons. The average represents the arithmetic average of the frequencies of the six species in each length bin.

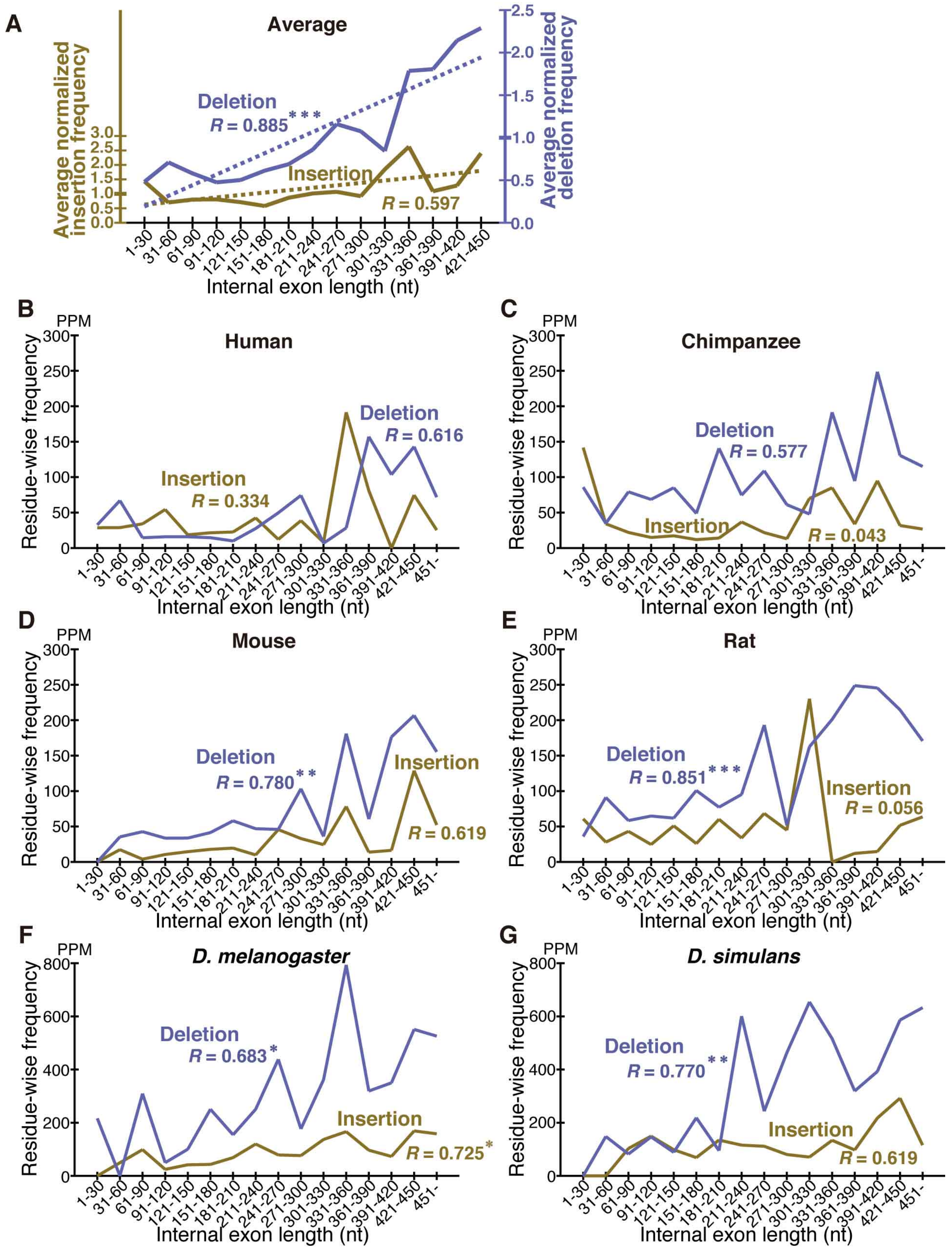


**Supplemental Figure S6.** Dependence of the frequency of the permissive sets of fixed indels on the length of internal exons. Legend as in Fig. 5.

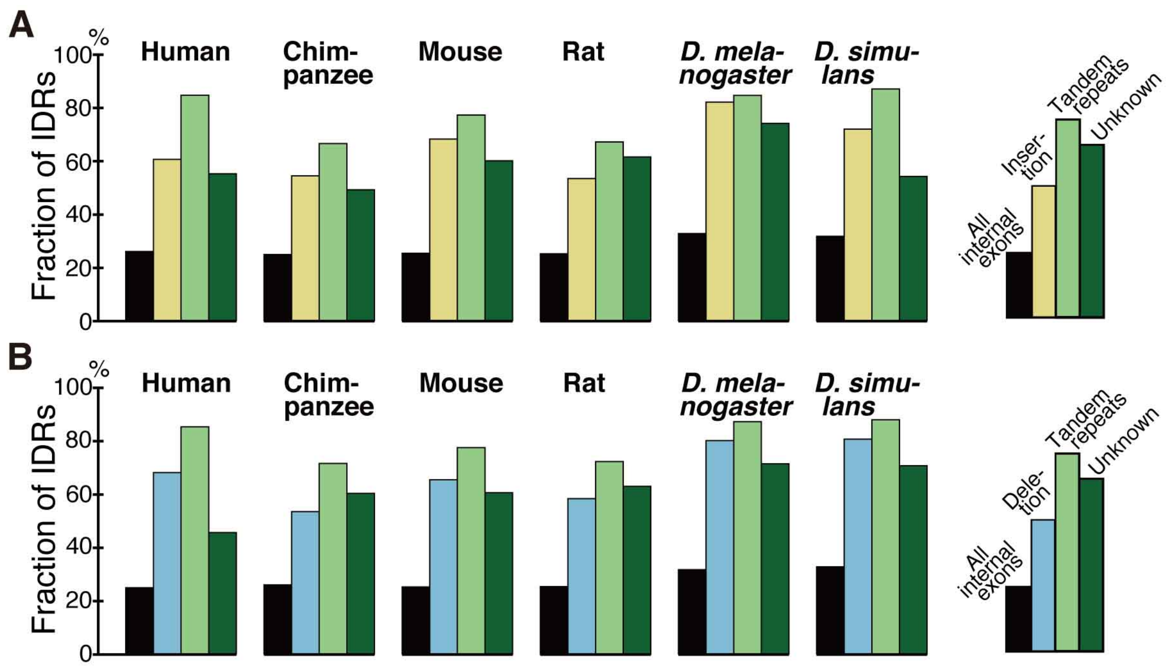


**Supplemental Figure S7.** Indels mostly encode IDRs in all species.

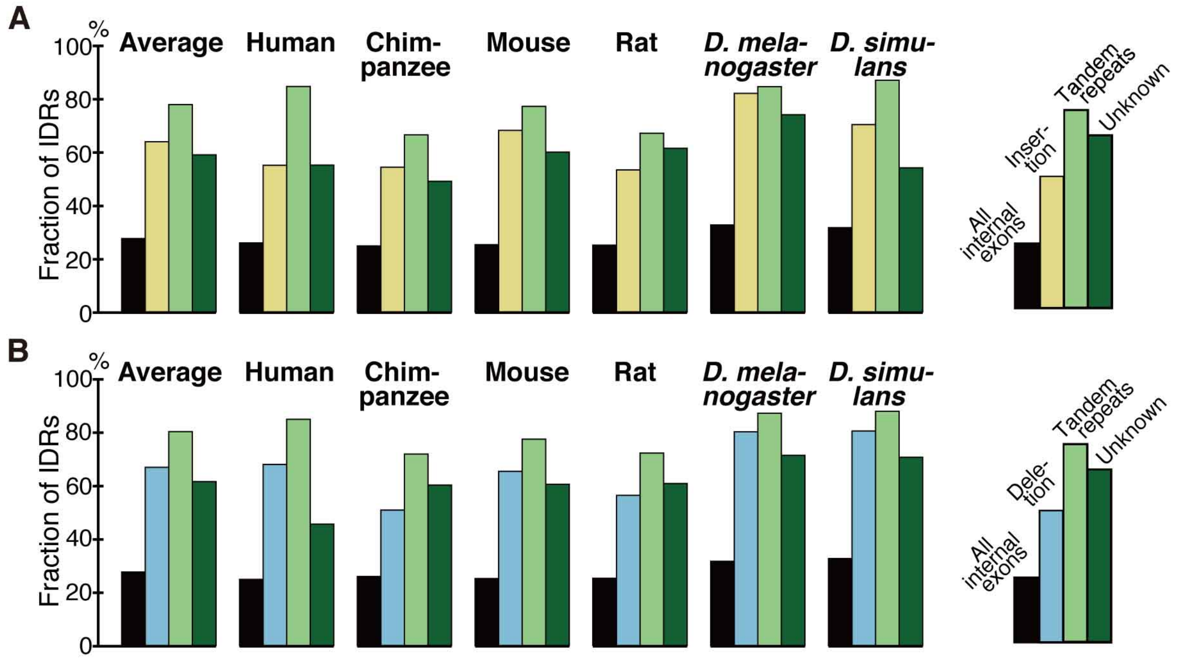


**Supplemental Figure S8.** Fixed indels of the permissive sets mostly encode IDRs.

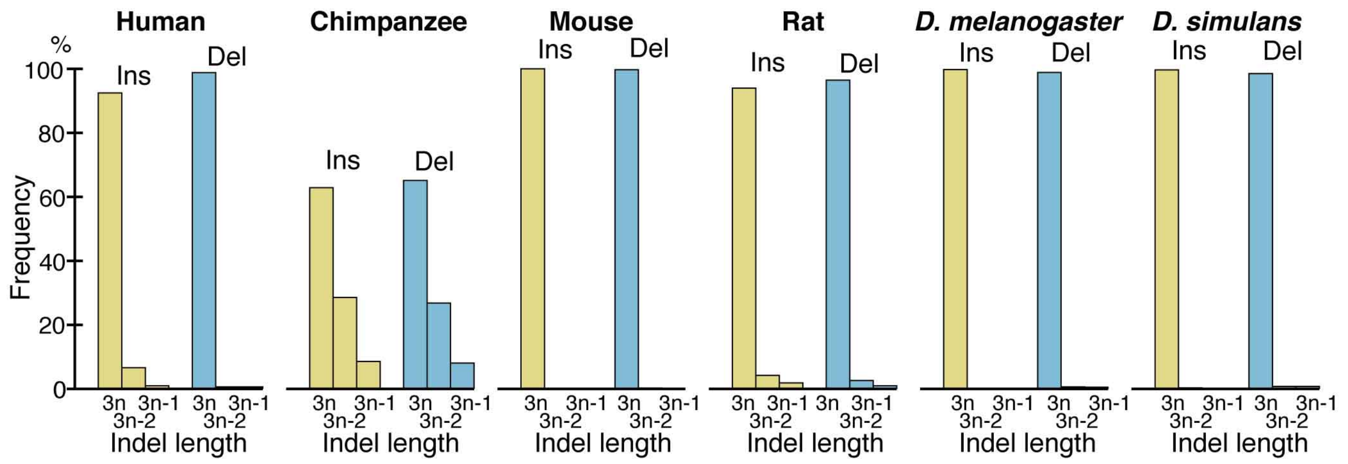


**Supplemental Figure S9.** Nearly all fixed indels preserve the reading frame.

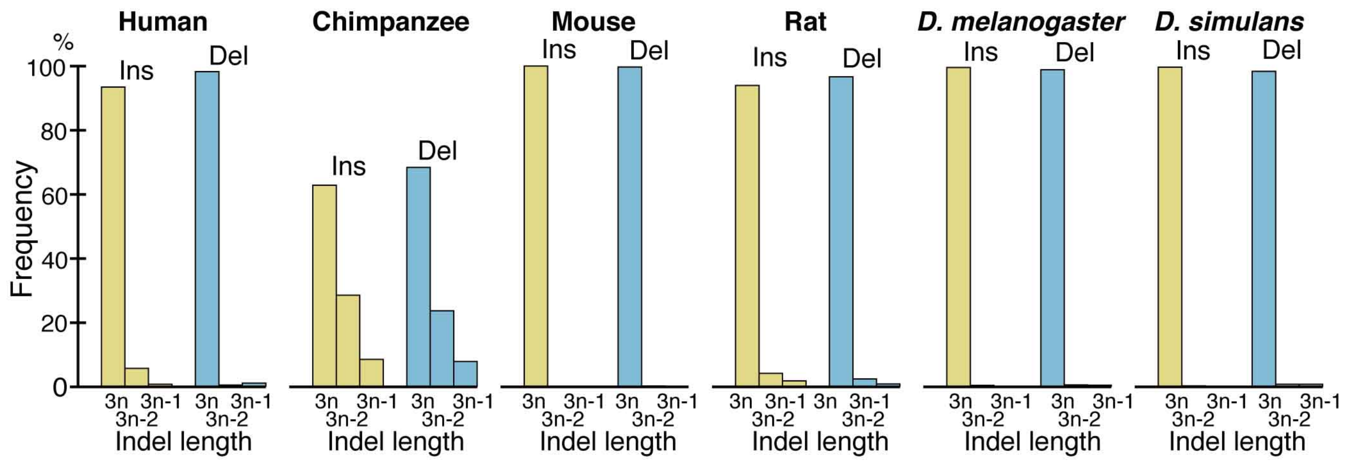


**Supplemental Figure S10.** Almost all fixed indels of the permissive sets preserve the reading frame.

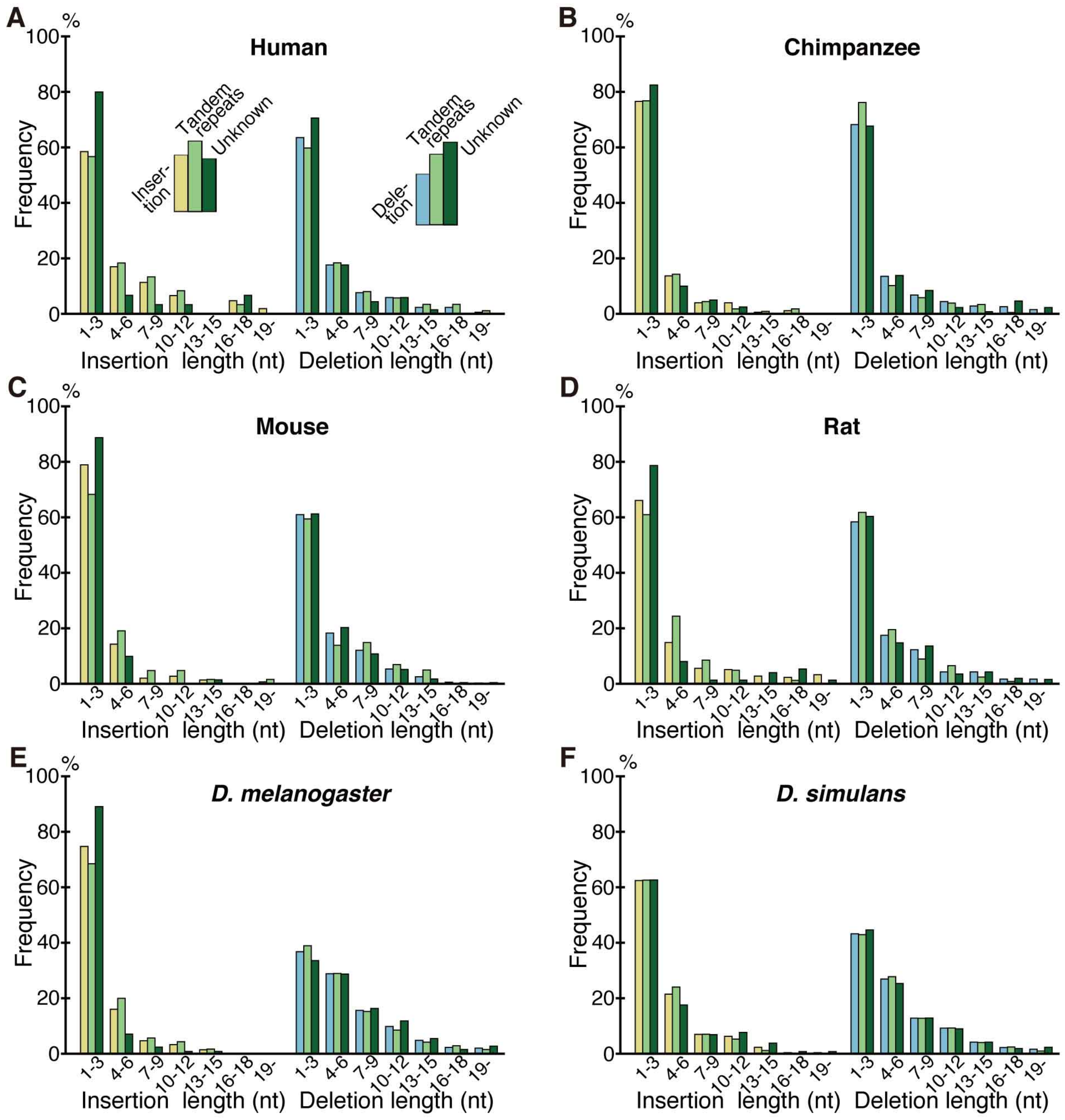


**Supplemental Figure S11.** Length distributions of fixed indels in each species.

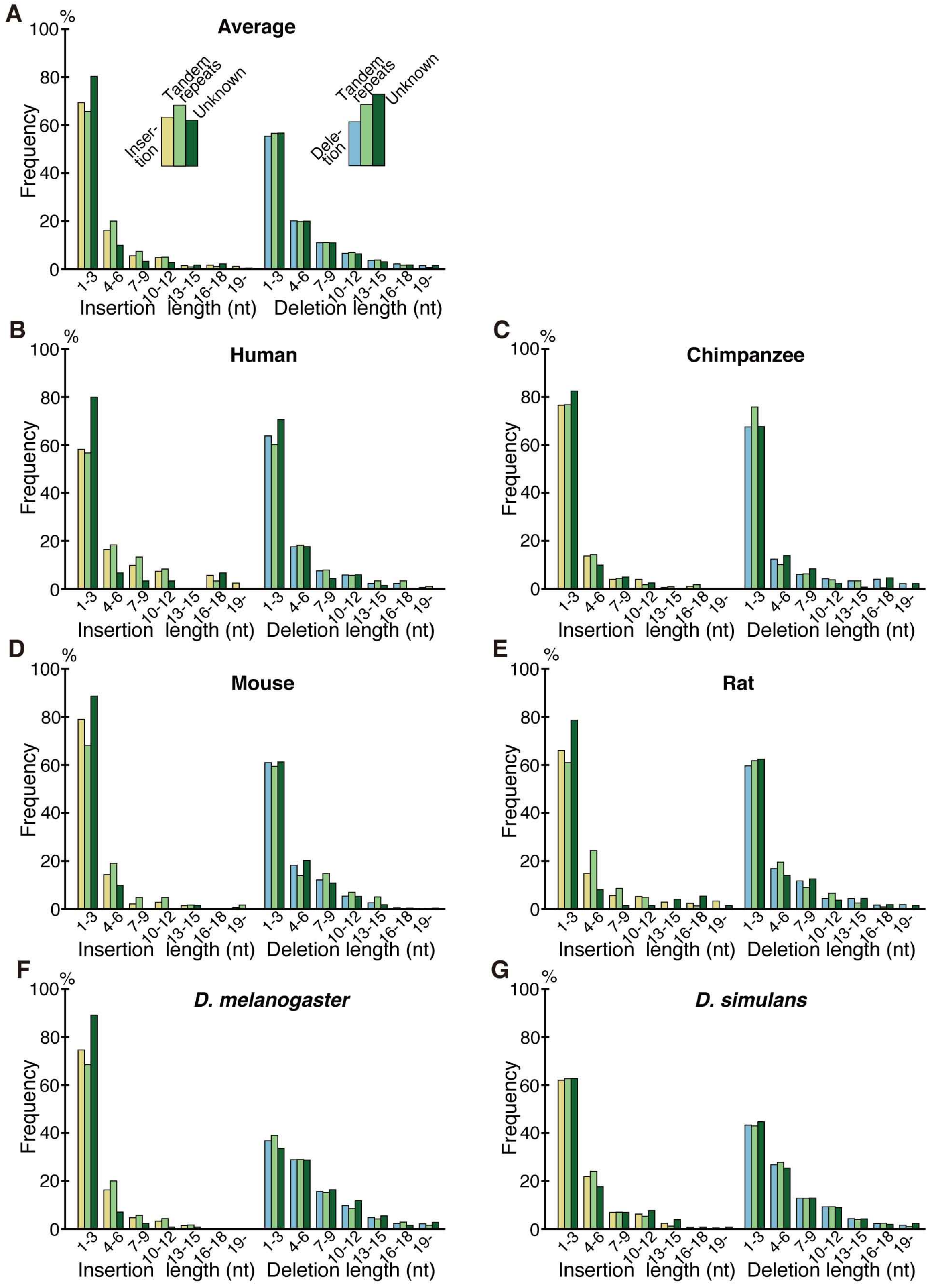


**Supplemental Figure S12.** Length distributions of the permissive sets of fixed indels.

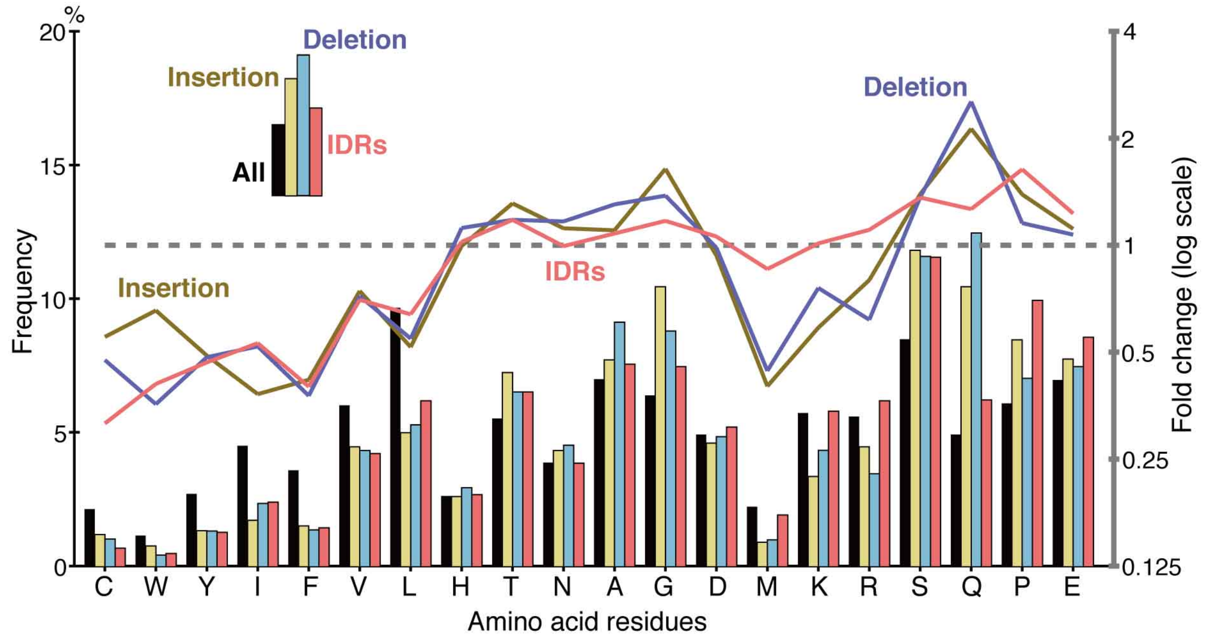


**Supplemental Figure S13.** Amino acid compositions of the permissive sets of fixed indels resemble those of IDRs. Legend as in Fig. 6*D*.

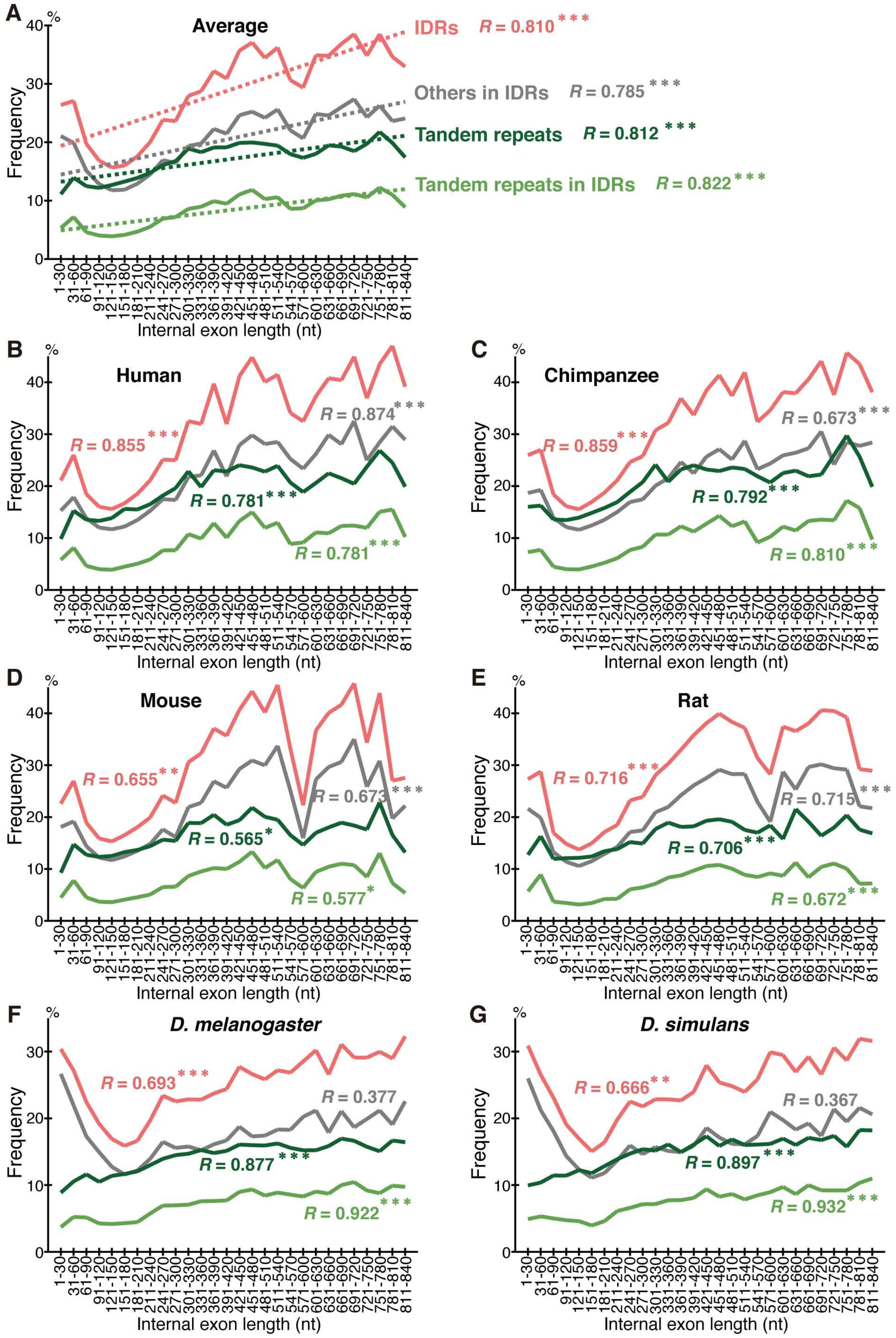


**Supplemental Figure S14.** Long internal exons tend to have a high fraction of repeats as well as IDRs. Legend as in Fig. 7.

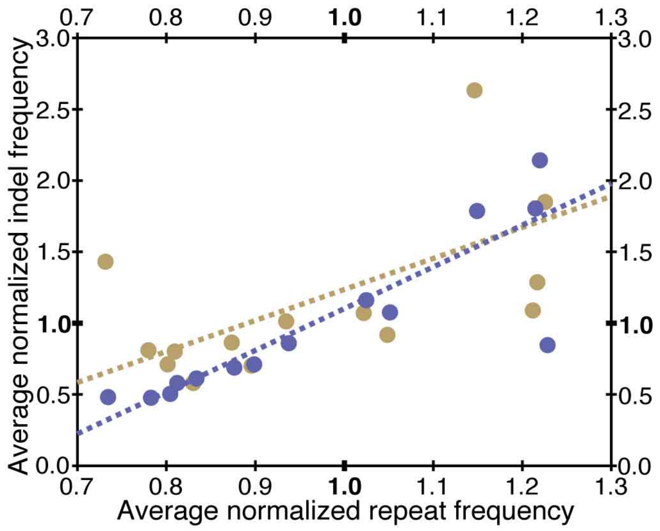


**Supplemental Figure S15.** Indel frequencies of the permissive sets and repeat frequency are correlated. Legend as in Fig. 8*A*.

-

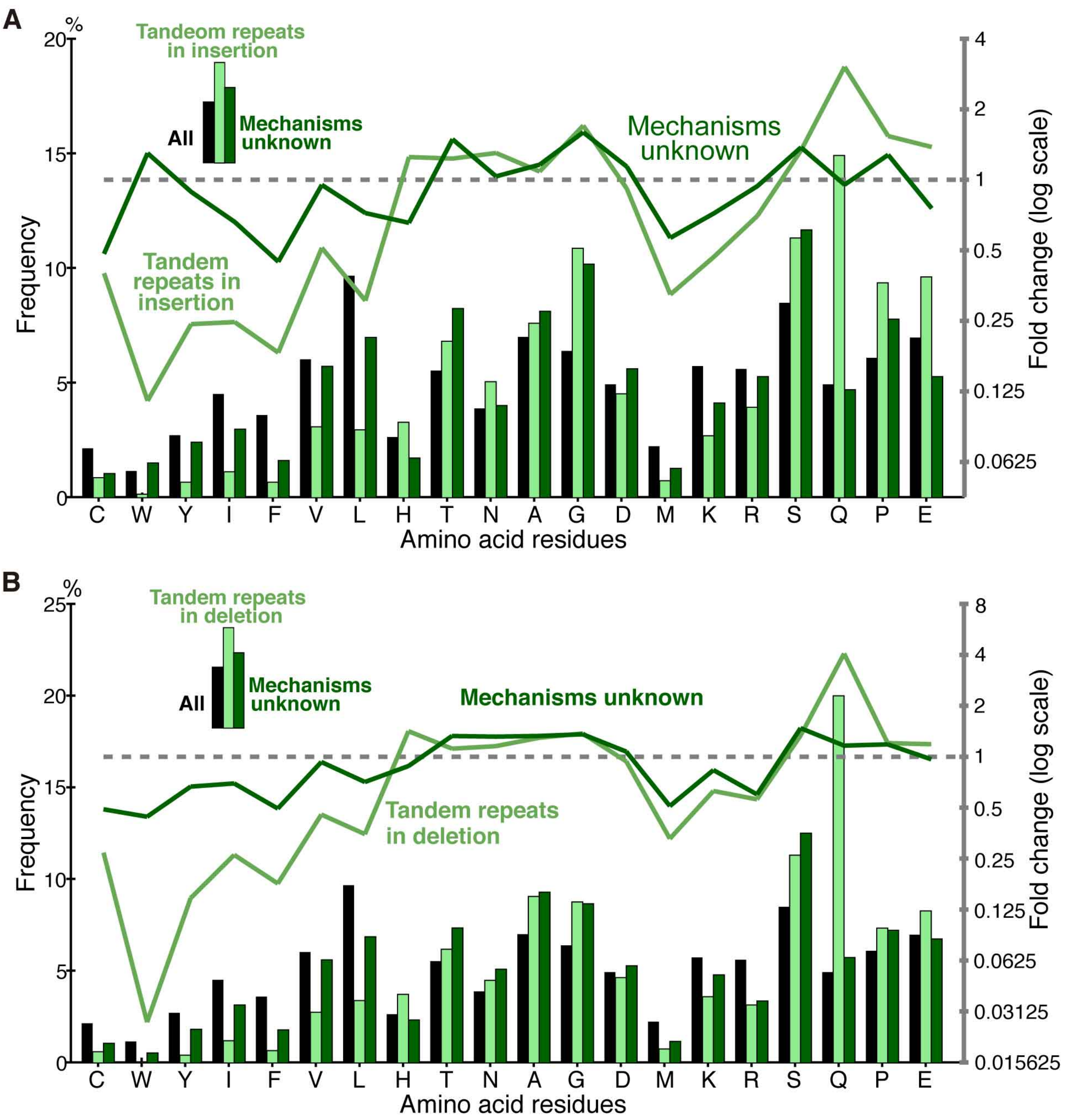


**Supplemental Figure S16.** Amino acid compositions of all internal exons, those generated by repeats, and those whose generation mechanisms are unknown. Legend as in Fig. 6*D*.

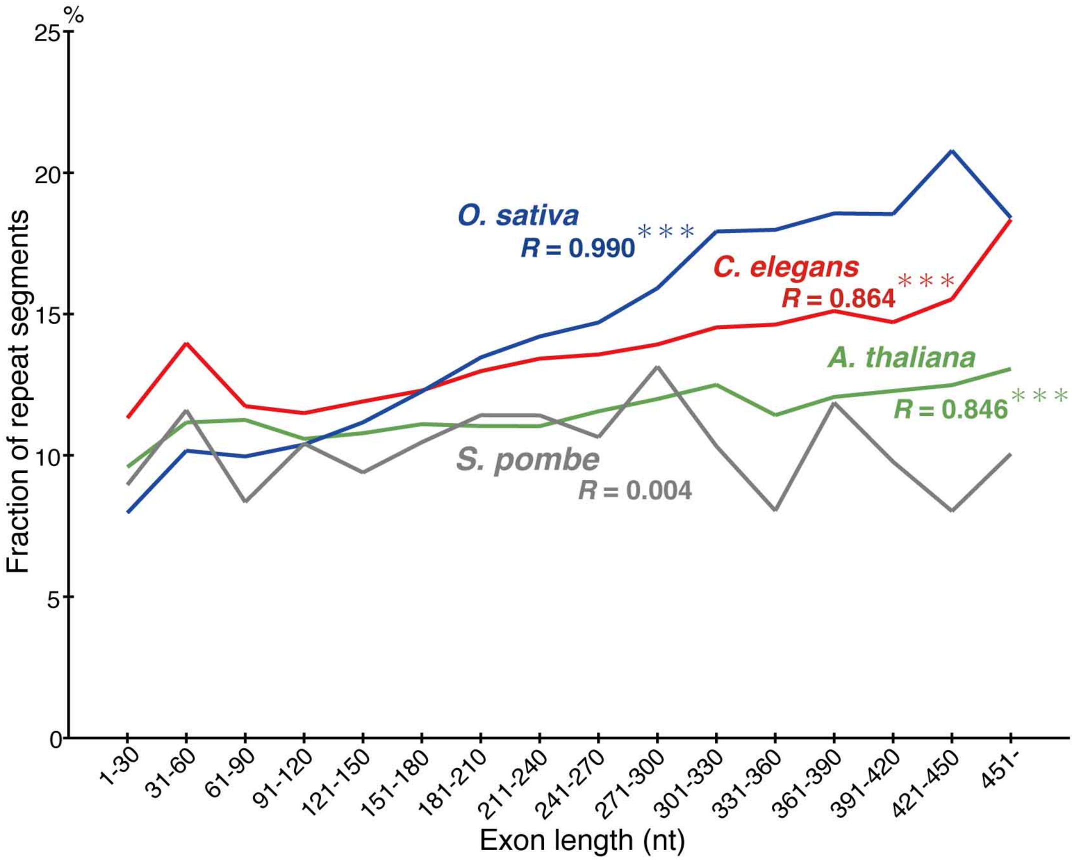


**Supplemental Figure S17** Long internal exons tend to have a high fraction of tandem repeats in *O. sativa*, *A. thaliana*, and *C. elegans*, but not in *S. pombe*. The fractions of tandem repeats are plotted with the correlation coefficients and statistical significance as in Fig. 5 legend.

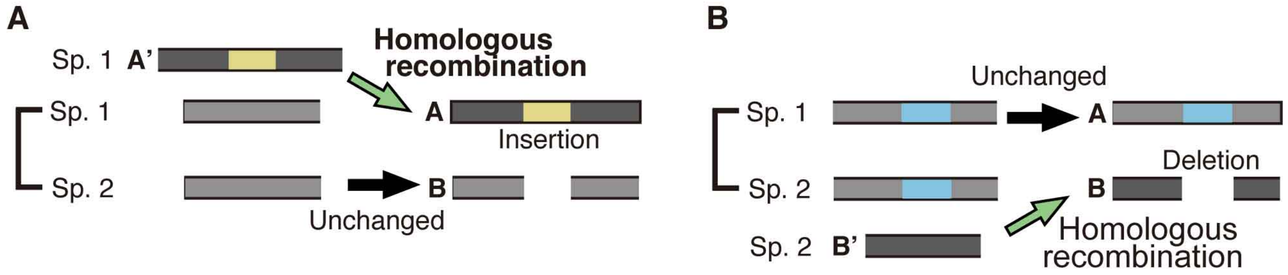


**Supplemental Figure S18.** Criteria for selecting indel cases generated by homologous recombination.
